# Spatial Transcriptomic Characterization of Novel Pathologic Niches in IPF

**DOI:** 10.1101/2023.12.13.571464

**Authors:** Christoph H. Mayr, Diana Santacruz, Sebastian Jarosch, Charlotte Lempp, Lavinia Neubert, Berenice Rath, Jan C. Kamp, Danny Jonigk, Mark Kühnel, Holger Schlueter, Jonas Doerr, Alec Dick, Fidel Ramirez, Matthew J. Thomas

**Author notes:** Correspondence should be addressed to: Matthew J. Thomas or Fidel Ramirez.

## Abstract

An unmet medical need persists in Idiopathic Pulmonary fibrosis (IPF), for which treatments additional to anti-fibrotic therapy are needed. Single cell RNA sequencing (scRNA-seq) has advanced our understanding of IPF with cell type-specific insights but lacks cellular tissue context. Spatial transcriptomics addresses this by providing spatially resolved gene expression, enabling gene and cell type localization within the tissue environment. We profiled IPF and control patient lung tissue sections using spatial transcriptomics and combined the data with an atlas of integrated IPF scRNA-seq datasets. Through computational analysis, we identified three disease-associated pathologic niches with unique cellular composition / localization and analyzed their cell-cell communication. We identified the Fibrotic niche, comprising Myofibroblasts and Aberrant Basaloid cells, preferentially located around airways and close to the Airway Macrophage niche in the lumen, containing SPP1+ Macrophages. We also identified the Immune niche, distinct foci of lymphoid cells in fibrotic tissue, surrounded by remodeled endothelial vessels.

**TEASER:** Spatial transcriptomics localizes genes and cell types in the tissue and identifies pathological cellular niches in IPF and control lungs.

## INTRODUCTION

Recent studies have harnessed the power of single-cell RNA sequencing (scRNA-seq) to enable a cell type-specific viewpoint on pathological changes in lung diseases, which rank third for causes of human mortality(*1–3*). Novel fibrosis-specific cell types and molecular changes associated with idiopathic pulmonary fibrosis (IPF) have been reported in such scRNA-seq studies, building a basis for a new understanding of fundamental biological processes occurring in the diseased fibrotic lung(*4–9*). Despite the availability of two anti-fibrotic therapies for IPF(*10*, *11*), a profound unmet medical need persists, requiring a next wave of therapies which address novel pathobiology beyond myofibroblast activity. The position of a cell, its surrounding neighbors and tissue structures, such as honeycomb morphology or fibroblastic foci in IPF, provide crucial information for defining cellular phenotype, disease state, and ultimately cell function and communication(*12*, *13*). During the sampling for scRNA-seq, however, the original tissue structure is inevitably destroyed, and the single cells have lost the spatial location information of their tissue context(*14*). In addition, the biased mechanical extraction process can result in the loss of more fragile cell types (e.g., alveolar type 1 (AT1) cells), thus distorting cell population distribution(*15*). Therefore, an unbiased understanding that goes beyond single marker imaging, of where fibrotic cell types such as Myofibroblasts, Krt5-/Krt17+ Aberrant basaloid cells, SPP1+ Macrophages or PVLAP+ Bronchial Vessels are located and the characteristics of their cellular neighborhood in the IPF lung, is lacking(*5–9*, *16*).

Spatial transcriptomics addresses this by providing spatially resolved gene expression within the natural tissue environment, of intact tissue without inducing cell stress, death, or an isolation bias(*13*, *17*, *18*). The recently established Visium for FFPE technology from 10X Genomics, enables an unbiased probe-based near whole-transcriptome wide mRNA analysis of preserved FFPE tissue sections up to an area of 11mm by 11mm(*14*, *19*).

Here, we use this technology to investigate the spatial transcriptomic profiles of IPF and control patient FFPE tissue sections. We also generated an integrated pulmonary fibrosis and interstitial lung disease (PF-ILD) scRNAseq atlas from published datasets. Combining those two modalities, we used the scRNA-seq data to deconvolute the cell type composition of Visium spots to map well defined cell types and genes onto the tissue. Applying bioinformatic analysis, we discovered eight distinct cellular tissue microenvironments, or niches, with unique cellular composition and localization, three of which were unique to IPF tissue: the Fibrotic niche composed by Myofibroblasts and Aberrant Basaloid cells, the Airway Macrophage niche containing SPP1+ Macrophages, and the Immune niche, foci with mostly lymphoid cells, surrounded by deranged endothelial cells. Importantly, we could use the spatial cell type information to rebuild the niche within the scRNA-seq data for a spatially educated and in-depth cell-cell communication analysis. Thus, we were able to reveal detailed tissue-based pathologic cellular crosstalk providing a resource for drug discovery aimed at disrupting disease-driving cellular niches in IPF.

## RESULTS

### Multi-omic profiling of healthy and IPF lung tissue

To localize IPF disease associated cell types in situ in the tissue context and gain insight into their cellular neighborhood, we performed spatial transcriptomics using the 10x Genomics Visium for FFPE and Visium CytAssist for FFPE platforms (Fig. 1a). We profiled 11 H&E-stained tissue sections from three IPF and three control patients (Supp. Fig. 1a and Supplementary table S2). Spots from all slides were integrated into one data manifold, representing 57,787 spots with 12,486 consistently expressed genes, after sample processing and quality control (QC) (Supp. Fig. 1b). We assessed the spatial distribution of cells in the tissue, by estimating the cell-type compositions of each spot, thereby increasing its resolution.

**Figure 1:**
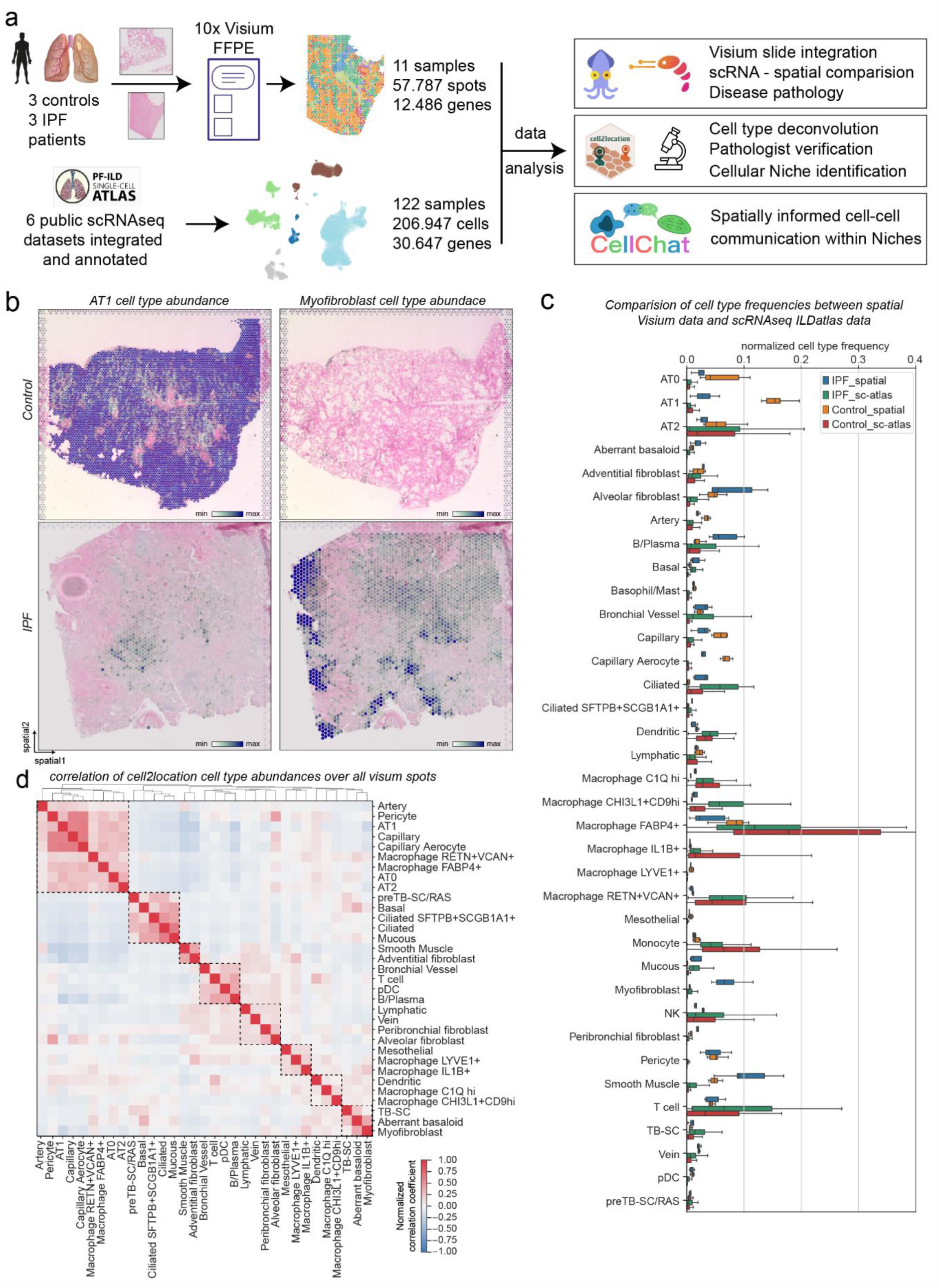
Combining spatial transcriptomics and scRNAseq places genes and cell types to their tissue location. **(a)** Experimental design included spatial transcriptomics using 10x Visium for FFPE and CytAssist on control and IPF patient samples (6 donors, 4x sections Visium CytAssist for FFPE, 7x sections with Visium for FFPE, see Supp. Fig. 3a). Six publicly available scRNAseq datasets were integrated and annotated into an PF-ILD atlas. Visium and scRNAseq data was combined for various analysis steps, including cell type deconvolution or cell-cell communication. **(b)** Spatial plots show cell2location mapping of estimated cell type abundance of AT1 and Myofibroblast cells on one Control and one IPF tissue section. **(c)** Comparison of normalized cell type frequencies in spatial transcriptomic data and scRNAseq IPF-ILD atlas. **(d)** Pearson correlation of cell type abundances estimated with cell2location across all tissue sections identifies spatial co-located cell type modules.

Therefore, we integrated six previously published scRNA-seq datasets(*4*, *5*, *7*, *8*, *20*, *21*) of interstitial lung diseases (ILD) and control tissue from human lungs into a pulmonary fibrosis – interstitial lung disease (PF-ILD) atlas to ensure reproducibility and increase statistical power (as described in Materials and Methods) (Fig. 1a). For this study, focusing on IPF and non-fibrotic controls, our PF-ILD atlas represents gene expression profiles of 206,947 single-cells from 122 human individuals (IPF n = 65, controls n = 57) across all six studies (Supp. Fig. 2b). The final annotation based on established single-cell signatures in the human lung(*9*) (Supp. Fig. 2c-j and Supp. Fig. 3a-d) represents 37 cell populations, characterized by distinct marker gene expression profiles, including recently reported pathologically preserved SPP1+ Macrophages, CTHRC1+ Myofibroblasts, KRT5-/KRT17+ Aberrant Basaloid cells and PVLAP+ Bronchial Vessels (Supp. Fig. 3e, f and Supplementary table S1).

Using the cell2location software(*22*), each spot was deconvoluted based on the annotated scRNA-seq data of the integrated PF-ILD atlas (Fig. 1a). Comparison of histopathological annotation by a pathologist on the Hematoxylin and Eosin (H&E) stained tissue sections and estimated cell type abundance, confirmed successful mapping of well-described cell types to their expected location, such as Arteries and Veins or Smooth Muscle cells around large vessels, as well as Ciliated and Mucous cells to airways (Supp. Fig. 1 c, d, e). Furthermore, we could observe disease specific effects such as the decrease of AT1 cells and the emergence of *CTHRC1*+ Myofibroblasts in IPF tissue (Fig. 1b).

To evaluate how well cell type frequencies were represented in spatial transcriptomics data, we compared the Visium data with the integrated PF-ILD scRNA-seq atlas (Fig. 1c). While mesenchymal cell type or airway epithelial cell type frequencies were comparable between the modalities, myeloid cells such as all Macrophage subtypes were found with higher frequencies in scRNA-seq data. On the contrary, AT1 and Aberrant basaloid cells or Capillaries were better represented in the spatial data, validating the spatial transcriptomics approach for analysis of healthy and diseased lung tissue.

We correlated the estimated cell type abundances per spot with each other to account for the resolution of multiple cells per spot (Fig. 1d). The clustering revealed co-localized cell types, representative of histopathological regions such as Alveoli with AT1, alveolar type 2 cells (AT2), capillaries and pericytes, as well as Airways with Basal, Ciliated and Mucous cells. Additionally, the co-localized cell types indicated disease induced changes with Aberrant Basaloid and Myofibroblast clustering together or various immune cells like T- and B-cells together with Bronchial Vessels.

In summary, the combination of an integrated scRNA-seq PF-ILD atlas together with spatial transcriptomics offers a new perspective on the changes in IPF and enables localization and analysis of discrete cell types in their tissue context.

### Three distinct cellular niches appear only in IPF lung tissue

To explore the spatial organization of lung tissue, we performed non-negative matrix factorization (NMF) using the cell2location spot-by-cell-type output. Hypothesizing that cellular niches could serve as structural and functional building blocks composed of characteristic cell types, we used NMF to identify them in an unbiased manner across all sections and conditions (Fig. 2a). We then evaluated factors of co-occurring cell types (Supp. Fig. 4a) and identified eight cellular niches with distinct cell type profiles and corresponding marker gene signatures (Supp. Fig. 4b and Supplementary table S3).

**Figure 2:**
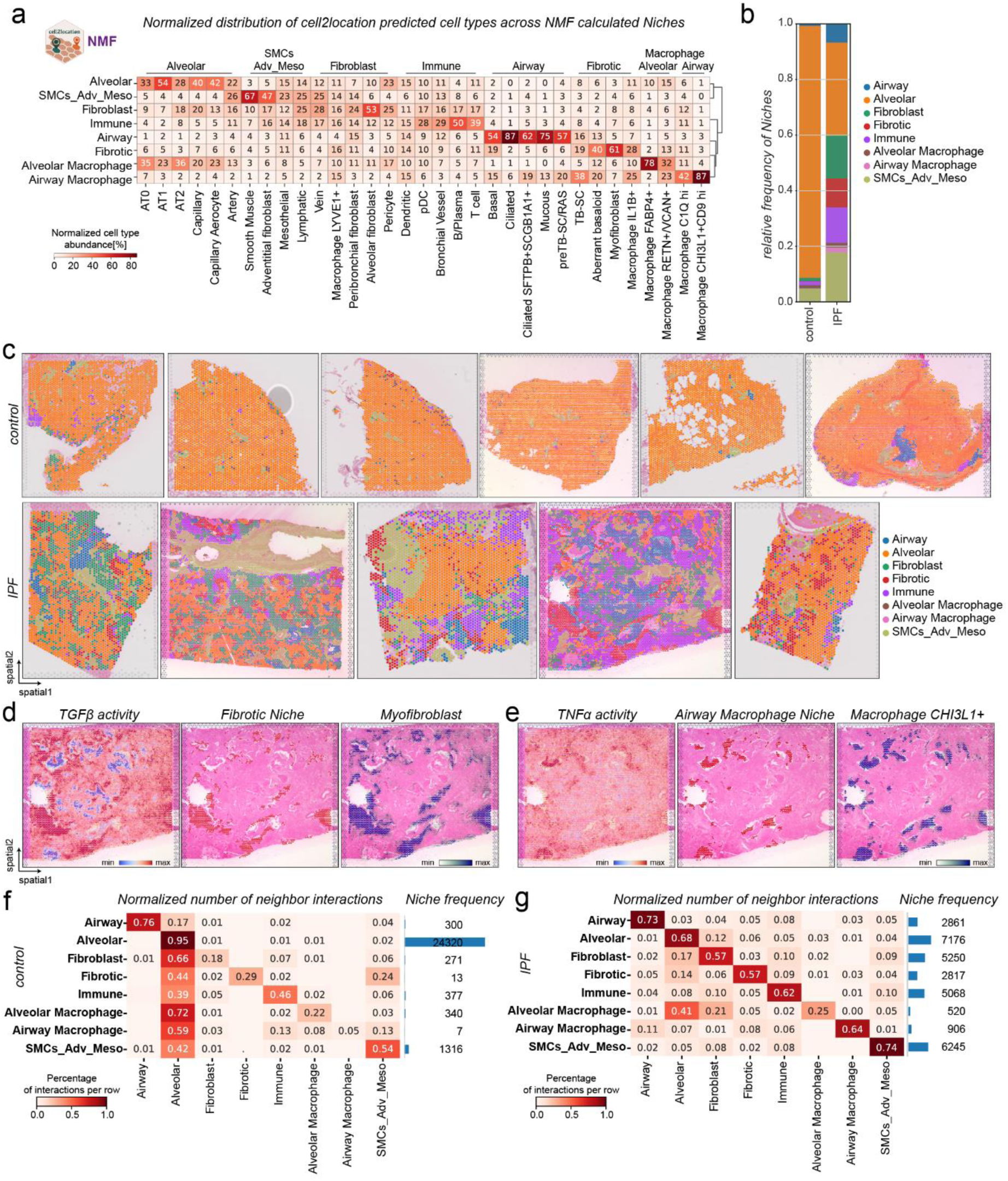
identification of three characteristic cellular niches in IPF lung tissue. **(a)** Heatmap shows the normalized distribution of estimated cell type abundance in percentages across the niches, that were computed using the NMF method of the cell2location package. SMCs_Adv_Meso refers to Smooth muscle, Adventitial fibroblast, and Mesothelial cells. **(b)** Relative frequency of niches across disease conditions is shown. **(c)** Spatial plots visualize the niche annotation across all samples and across conditions. **(d-e)** Spatial plots show characterization of spatial transcriptomics data by pathway activity, niche annotation and cell types for one IPF section. **(f-g)** Heatmaps show normalized neighbor interactions for each Niche in **(f)** control and **(g)** IPF tissue.

We next assessed the relative frequency of these niches in control and IPF tissue. While control tissues were dominated by the Alveolar niche, containing mainly AT1, AT2 and Capillaries, the Airway niche, the Fibroblast niche, the Smooth Muscle/Adventitial/Mesothelial niche, as well as the Alveolar Macrophage niche could still be found in small quantities (Fig. 2b). The Airway niche contained the highest portion of all airway epithelial cells with Basal, Ciliated, Ciliated *SFPTB*+*SCGB1A*+, Mucous and pre-Terminal Bronchial Secretory cells (pre-TB-SCs).

The Fibroblast niche contained the highest proportion of different fibroblast subsets, while the mixed Smooth Muscle/Adventitial/Mesothelial niche contained several structural cell types such as Smooth Muscle Cells (SMCs), Arteries, Adventitial fibroblast or Mesothelial cells. Interestingly, the Alveolar Macrophage niche, while containing the highest proportion of FAPB4+ Macrophages as well as RENT+/VCAN+ Macrophages, still contained decreased numbers of alveolar subtypes such as AT1 cells or capillaries, but the highest proportion of AT2 cells (Fig. 2a).

In IPF however, the Alveolar niche frequency decreased, and the frequency of all other niches increased (Supp. Fig. 4c), with three niches exclusively identified only in IPF tissue: (I) the Fibrotic niche containing Myofibroblasts, Aberrant Basaloid cells and surprisingly the airway-associated cell type known as Terminal Bronchial Secretory (TB-SCs)(*23*); (II) the Immune niche containing B- and T-cells, Dendritic cells and PVLAP+ ectopic endothelial Bronchial vessels; and (III) the Airway Macrophage niche containing SPP1+ Macrophage subsets of CHI3L1+CD9hi and C1Qhi Macrophages as well as TB-SCs (Fig. 1b, c and Supp. Fig. 4c and Supp. Fig. 5a).

To link cellular function to spatial cell composition within niches, we performed enrichment analysis on spatial niches. We used DGE to identify niche specific gene signatures between niches and did functional analysis using Molecular Signatures Database (MSigDB) as well as Gene Ontology (GO) and estimated signaling pathway activities (PROGENy and decoupleR)(*24*, *25*) for all spots in one niche. For example, the Fibrotic niche can be characterized with several ECM-associated GO-terms and MSigDB terms such as Epithelial Mesenchymal Transition or Hypoxia, and co-localized with areas of highest TGFβ signaling activity and Myofibroblast occurrence (Fig. 2d and Supp. Fig. 4d, e, f). Airway Macrophage niches were the areas of high TNFα signaling activity that colocalized with SPP1+ Macrophages and could be characterized with terms referring to Neutrophil activation, Interferon Gamma Response or IL6 regulation (Fig. 2e and Supp. Fig. 4d, e, f).

To explore the spatial organization of identified niches, we leveraged the positional data and analyzed neighboring interactions of the niches. The control tissue, as expected, was dominated by the Alveolar niche, being the most frequent neighbor for most of the niches except for the Smooth Muscle/Adventitial/Mesothelial niche, that especially with the Muscle regions captured on some sections could form bigger regions by itself (Fig. 2f). In IPF, the tissue looked unorganized, both when looking at the immediate (1-Ring) or extended neighborhood (3-Rings) of a spot (Supp. Fig. 4g). With the decrease of the Alveolar niche and the increase of all other niches in frequency, the niches mostly neighbored themselves, but also all niches interacted with each other (Fig. 2g). Interestingly, the Fibrotic niche, seemed to neighbor all other niches, especially Alveolar, Immune, Fibroblast and Airway niches (Fig. 2g).

In summary, the unbiased identification of cellular niches as functional tissue building blocks allowed the analysis of spatial organization across samples and the changes in disease, that revealed the presence of characteristic IPF tissue specific Fibrotic, Immune and Macrophage niches.

### Aberrant basaloid cells are found around fibrotic airways

To further study the Fibrotic niche, only appearing in IPF tissue, we first looked at the distribution across sections and confirmed the occurrence in all IPF tissue sections (Supp. Fig. 5a). Next, we focused on the cell types contained in the niche based on the NMF analysis (Fig. 2a) and verified co-localization of Myofibroblast, Aberrant Basaloid cells, TB-SC, Basal cells and IL1B+ Macrophages by way of example on two IPF tissue sections (Fig. 3a-d and Supp. Fig. 6a-c and 7a, c). Overlap of established cell type marker genes such as CTHRC1 and ASPN for Myofibroblast, MMP7 and KRT7 for Aberrant Basaloid cells, as well as SCGB3A2 for TB-SC, additionally validated this result (Supp. Fig. 7b, d, e). Surprisingly, we found the Fibrotic niche preferentially located around airways. Myofibroblasts could also be found alone in more alveolar parts, which agrees with previous reports that found IPF specific fibroblastic foci predominantly in the more damage-susceptible distal parenchyma(*27*, *28*). Aberrant basaloid cells visually localized to airways and were always found together with Myofibroblasts (Fig. 3b, d and Supp. Fig. 6a, b). Supporting this finding, Aberrant Basaloid cell abundance correlated with a fraction of Basal cells, as well as, with recently identified TB-SC cells that reside in the distinct location of terminal bronchi, an anatomical feature found in human but not in mice(*23*) (Fig. 1d, 2a and Supp. Fig. 4a). Aberrant basaloid cells have previously been associated with senescence pathways and genes(*5*, *6*, *29*, *30*). We therefore calculated a custom senescence gene score for every spot in the spatial data using the hotspot gene-module scoring(*26*), and visualized it on two IPF tissue sections, as well as across the niches in all sections. The score was highest in the Fibrotic niche and in the highlighted regions of Myofibroblast and Aberrant basaloid co-localization (Fig. 3e, f and Supp. Fig. 7e, f).

**Figure 3:**
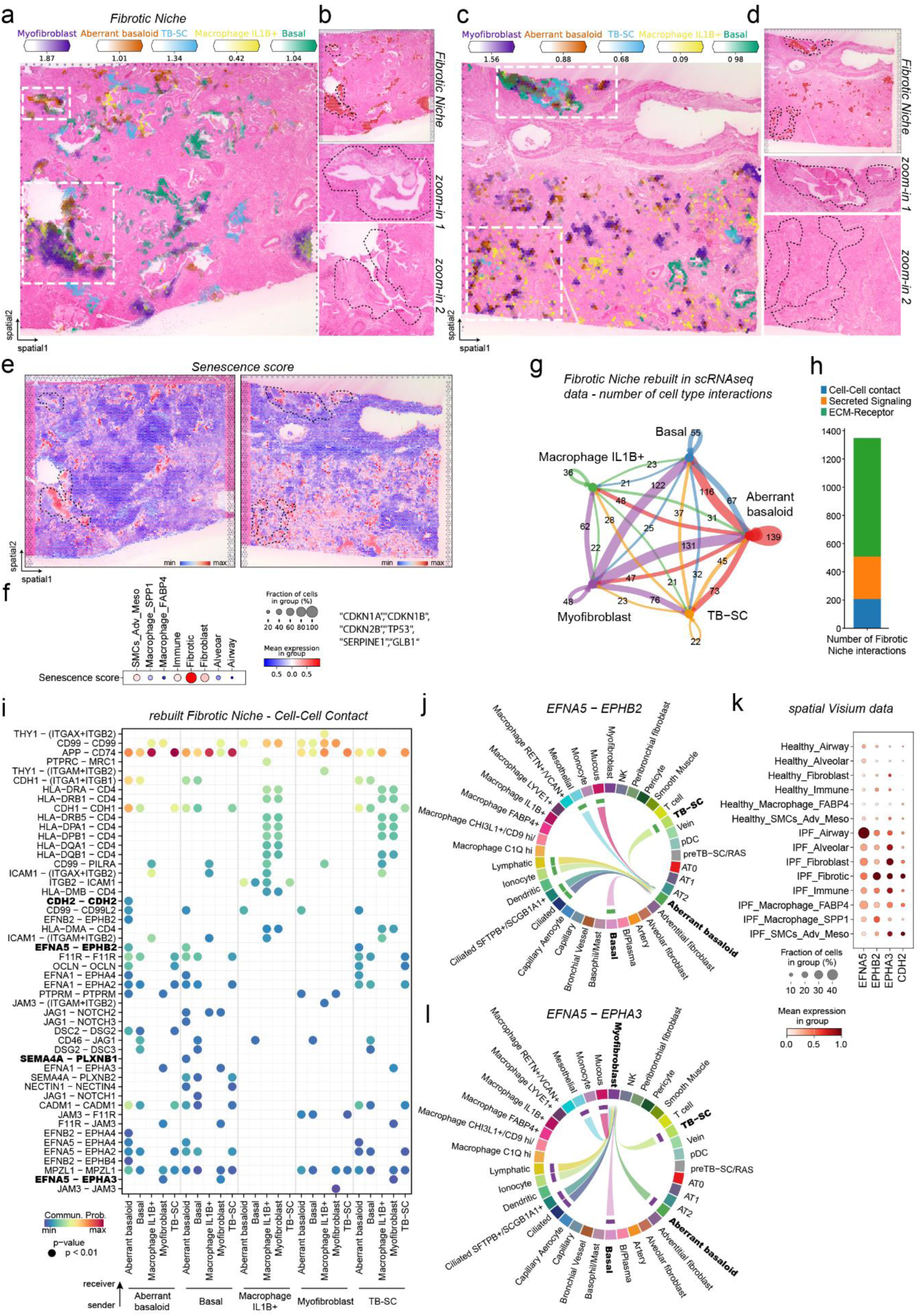
The Fibrotic niche localizes around airways. **(a-d)** Spatial plots show for two IPF tissue sections **(a, c)** abundance of Fibrotic niche associated cell types, **(b, d)** Fibrotic niche distribution as well as zoom into H&E-stained tissue for highlighted regions. **(e-f)** Senescence gene score (*CDKN1A*, *CDKN1B*, *CDKN2B*, *TP53*, *SERPINE1*, *GLB1*) calculated using hotspot gene-module scoring(***26***), **(e)** plotted on two IPF tissue sections, **(f)** plotted as dot plot against the other niches. **G)** Bubble plot summarizing the number of interactions between cell types within the rebuilt Fibrotic niche in the scRNA-seq PF-ILD atlas data. **(h)** Number of interactions within the Fibrotic niche per communication category. **(i)** Heatmap of significant ligand-receptor pairs from the Cell-Cell contact category within the Fibrotic niche. **(j-k)** Chord plot visualizing the interacting cell types for the ligand-receptor pair **(j)** *EFNA5*-*EPHB2* and **(k)** *EFNA5*-*EPHA3*. **(l)** Gene expression of given receptor and ligand genes across the niche and disease in the spatial Visium data.

To further understand the Fibrotic niche and the mechanism that regulated the dynamics of this cellular niche, we used CellChat(*31*) to perform cell-cell communication analysis. Making use of knowledge gained from the spatial Visium data, we used the estimated cell type abundances and rebuilt the Fibrotic niche in the higher resolution scRNA-seq data of the PF-ILD atlas (Supplementary table S4). Summarizing the number of all interactions revealed that Aberrant Basaloid cells had mostly interacted with Myofibroblast, directly followed by Basal cells and TB-SCs, providing further evidence for their airway like character (Fig. 3h). Cell-cell communication in the Fibrotic niche was dominated by over 60% of ECM-Receptor associated ligand-receptor pairs, that could be mostly assigned to Collagen, Laminin, Thrombospondin and Fibronectin pathways (Fig. 3g and Supp. Fig 8a). While Myofibroblast were responsible for most of these ECM interactions, also Aberrant Basaloid cells contributed with the expression of, for example Fibronectin (*FN1)* or Tenascin (*TNC*), consolidating their epithelial to mesenchymal transition (EMT) feature, unique among the epithelial cells (Supp. Fig. 9a-c). Yet, they were also able to receive ECM signals, for example from *TNC*, with the expression of the integrin receptors *ITGAV* and the characteristic *ITGB6*(*6*, *32*) (Sup. Fig. 9d).

In terms of secreted signaling, Midkine (MK) signaling was dominating the Fibrotic niche, followed by known fibrosis pathways such as Transforming Growth Factor Beta (TGFβ), Epithelial Growth Factor (EGF), Semaphorin (SEMA3) or Bone Morphogenic Proteins (BMP)(*33*) (Supp. Fig. 8a, b). That BMP signaling was having an influence on Aberrant basaloid cells became apparent with the interaction between Peribronchial and Alveolar fibroblast, as well as myofibroblast, with the ligand *BMP5* and the dimerizing receptor of *ACVR1* and *BMPR2*, only expressed by Aberrant basaloid cells (Supp. Fig. 8c).

Focusing on the Cell-Cell contact interactions, which we assumed could best be validated in the multiple cells containing Visium spots, we identified the Ephrin pathway as major contributor, besides the MHC-II pathways, Cadherin (CDH) or Notch signaling (Fig. 3i and Supp. Fig. 8a). The Ephrin pathway had been associated with proliferative disease such as cancer, but we found distinct interactions in the Fibrotic niche, suggesting a potential role in fibrosis(*34*). On the one hand, the receptor *EPHB2* is specifically expressed by Aberrant basaloid cells, on the other hand, the receptor *EPHA3* seemed to be specific for Myofibroblast. Both received signals from *EFNA5*, mostly expressed by airway epithelial cell types (Fig. 3i, j, k). Both observations could be validated in the spatial data, with *EPHA3* and *EPHB2* mostly expressed in the Fibrotic niche, while the ligand *EFNA5* is upregulated in IPF and mostly expressed in the Airway niche (Fig. 3k).

Overall, the combination of unbiased transcriptomics and tissue structure, revealed a preferential localization of the Fibrotic niche and especially Aberrant Basaloid cells around airways and allowed a spatially educated cell-cell communication analysis, that characterized signaling in fibrotic areas.

### SPP1+ Macrophage subsets localize to fibrotic airway lumen

To further study the Airway Macrophage niche, we first aimed to understand the heterogeneity of cellular Macrophage subtypes in IPF we first aimed to understand the heterogeneity of robustly identified cellular Macrophage subtypes in IPF, that all shared expression of Osteopontin (*SPP1*), recognized as being expressed by Macrophage across fibrotic diseases and organs(*7*, *35*). CHIT3l1+ Macrophages, resembling previously identified TREM2+ Macrophages(*36*), defining the Airway Macrophage niche (Fig. 2a), and C1Qhi Macrophages(*36*), both were nearly exclusively found in IPF tissue (Supp. Fig. 10a, b).

C1Q Macrophages, while contributing 42% to the Airway Macrophage niche, also localized to other niches such as the Fibroblast or Immune niche and were therefore found throughout the tissue (Fig. 2a, Supp. Fig. 10b). IL1B+ Macrophages co-localized with both the Fibrotic niche as well as the Airway Macrophage niche in distinct histopathological regions (Sup. Fig. 10c).

Next, we focused on the cell types contained in the niche based on the NMF analysis (Fig. 2a) and verified co-localization of CHIT3l1+ and C1Qhi Macrophages together with Aberrant Basaloid cells, TB-SC, SFTPB+ Ciliated cells and preTB-SC/RAS cells on IPF tissue sections (Fig. 4a-d and Supp. Fig. 11a, c). Overlap of established cell type marker genes such as *CHI3L1* and *CHIT*, or *C1QA* and *CCL13* for Macrophages, as well as *CAPS*, *SFTPB* and *SCGB1A1* for the Ciliated cells, additionally validated this result (Supp. Fig. 11b, d, e). We uncovered that the airway Macrophage niche was preferentially located to the distal airway lumen. *SFTPB* had recently been identified to do discriminate distal airway epithelium from proximal cell populations(*23*). That supports our finding with the Airway Macrophage niche colocalizing with SFTPB+ Ciliated cells but not SFTPB-Ciliated cells, in what seem to be rather large but distal airways (Fig. 2a and Fig. 4a and Sup. Fig. 11). Having identified a similar location for the Fibrotic niche, we looked at the distribution of both niches across the IPF sections. The Airway Macrophage niche, located in the lumen, neighbored the Fibrotic niche, located around the same airway, in some large airways while the Fibrotic niche could additionally be found in more parenchymal tissue areas (Supp. Fig. 11f). Checking the cell type distribution across the niches, Myofibroblast were rarely found outside the Fibrotic niche, but 20% of the Aberrant basaloid cells also contributed to the Airway Macrophage niche, verified by co-localization (Fig. 2a and Fig. 4a, c).

**Figure 4:**
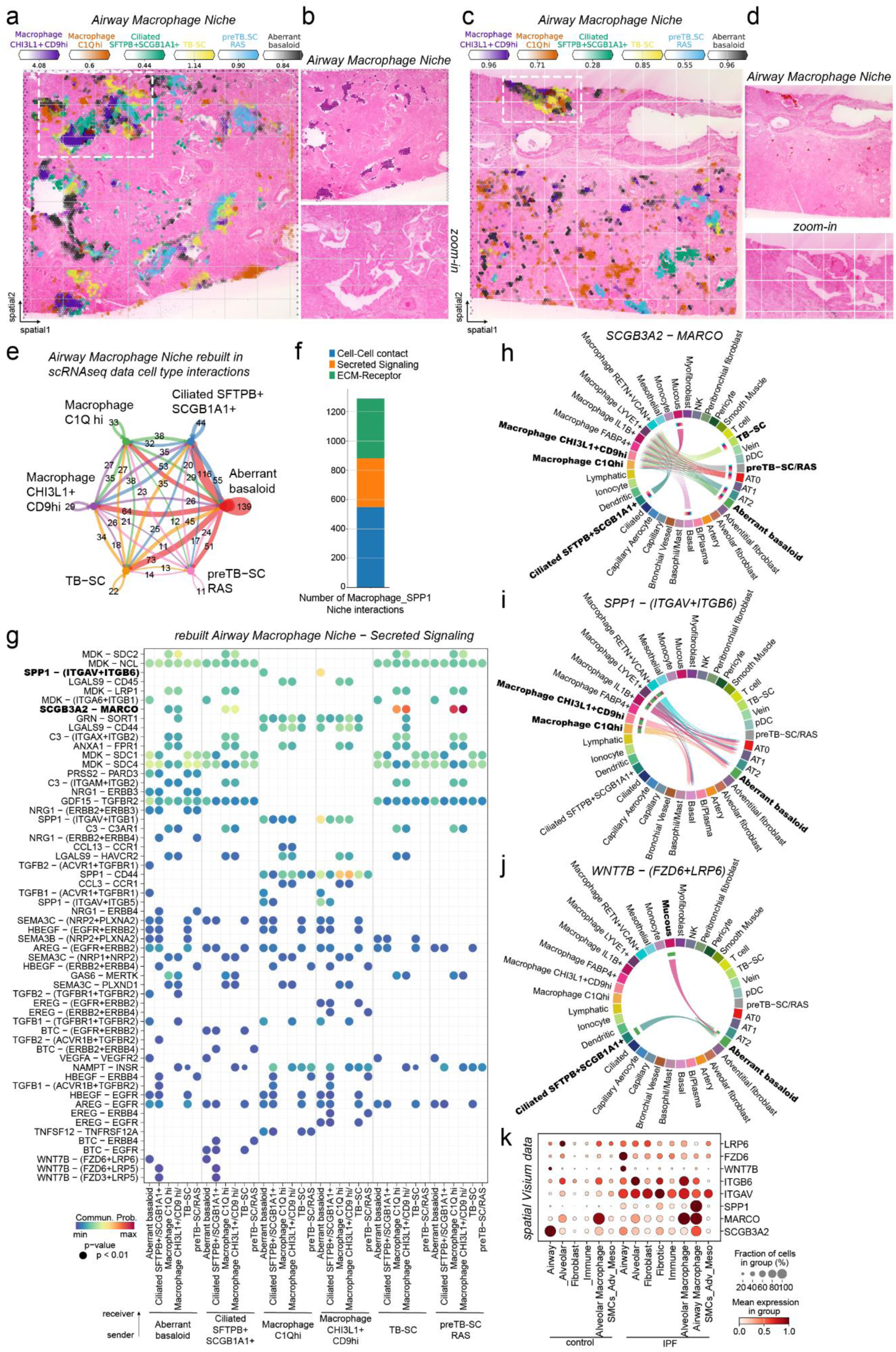
SPP1+ Macrophages are localize to fibrotic airway lumen. **(a-d)** Spatial plots show for two IPF tissue sections **(a, c)** abundance of Airway Macrophage niche associated cell types, **(b, c)** niche distribution as well as zoom into H&E-stained tissue for highlighted regions. **(e)** Bubble plot summarizing the number of interactions between cell types within the rebuilt Airway Macrophage niche in the scRNA-seq PF-ILD atlas data. **(f)** Number of interactions within the Airway Macrophage niche per communication category. **(g)** Heatmap of significant ligand-receptor pairs from the Secreted Signaling category within the Airway Macrophage niche. **(h-j)** Chord plot visualizing the interacting cell types for the ligand-receptor pair **(h)** *SCGB3A2*-*MARCO*, **(i)** *SPP1*-(*ITGAV*+*ITGB6*) and **(j)** *WNT7B*-(*FZD6*+*LRP6*). **(k)** Gene expression of given receptor and ligand genes across the niche and disease in the spatial Visium data.

To further analyze the Airway Macrophage niche, we rebuilt it again with its contributing cell types in the scRNA-seq dataset and performed analysis of the cell communication (Supplementary table S5). Summarizing the number of all interactions revealed that Aberrant Basaloid with their mixed epithelial and mesenchymal character were showing the most interactions within the niche, with SFTPB+ Ciliated cells, TB-SC and CHI3L1+ Macrophages (Fig. 4e). Cell-cell communication in the Airway Macrophage niche could mostly be assigned to Cell-Cell contact interactions (44%), followed by ECM-Receptor (Supp. Fig. 13a) and secreted signaling interactions (Supp. Fig. 12b), in contrast to the Fibrotic niche (Fig. 4f). Most of the Cell-Cell Contact associated ligand-receptor pairs, could be assigned to the MHC-II pathway (Supp. Fig 12a). Here the interaction of for example *HLA-DMB* expressed by all the Macrophage subtypes was shown to interact with *CD4*, expressed on Ciliated cells and Myofibroblasts, supporting the airway localization of the Airway Macrophage niche (Supp. Fi. 12b, c).

The interactions in the Secreted Signaling also offer possible explanations for the airway localization. Secretoglobin family 3A member 2 (*SCGB3A2*) expressed by all the airway epithelial cell types as well as Aberrant basaloid cells, binds to the Macrophage Receptor With Collagenous Structure (*MARCO*), expressed by all macrophages (Fig.4g, h). Also, the ligand *SPP1*, can bind to dimerizing pair of integrins *ITGAV* and *ITGB6*, expressed by Aberrant Basaloid cells, suggesting a distinct communication between these two cell types only arising in fibrotic conditions (Fig. 4i). Aberrant basaloid cells seemed further stimulated by specific WNT signaling, secreted by Mucous and SFTPB+ Ciliated cells in the form of WNT7B, binding to the dimerizing receptor of *FZD6* and *LRP6* (Fig. 4j). All three ligand-receptor observations could be validated in the spatial data, with *SCGB3A2* and *SPP1* mostly expressed in the Airway Macrophage niche in IPF, while *MACRO* and the integrins *ITGAV* and *ITGB6*, as well as the WNT receptors *FZD6* and *LRP6* are upregulated in IPF and expressed in the Airway Macrophage and Airway niche, respectively (Fig. 3k).

Overall, our spatial data revealed the differential localization of Macrophage subtypes in IPF and suggested a preferential localization of the Airway Macrophage niche to the lumen of fibrotic airways, in close proximity the Fibrotic niche and especially Aberrant basaloid cells, via distinct cell-cell communication.

### Immune cell foci are recruited by IPF-specific ectopic endothelial Bronchial Vessels

The third pathologic niche appearing in all IPF tissue sections was the Immune niche, consisting of B- and Plasma cells, T cells, Plasmacytoid dendritic cells (pDCs) and the recently discovered *PLVAP*+ ectopic endothelial cell population here named Bronchial Vessels(*16*, *37*, *38*) (Supp. Fig. 5b). While all cell types contributing to the Immune niche show a dramatic increase in IPF tissue compared to controls (Supp. Fig. 14), we noticed them clustering together in distinct foci, even visible in the H&E staining by a grey color (Fig. 5a, b). We verified co-localization of the contributing cell types in IPF tissue sections (Fig. 5a, b and Supp. Fig. 15a, c), as well as overlap of established cell type marker genes such as *CD19*, *CD79A* and *MS4A1* for B- and Plasma cells, *CD3D* for T cells, as well as *PLVAP* for Bronchial Vessels (Supp. Fig. 15b).

**Figure 5:**
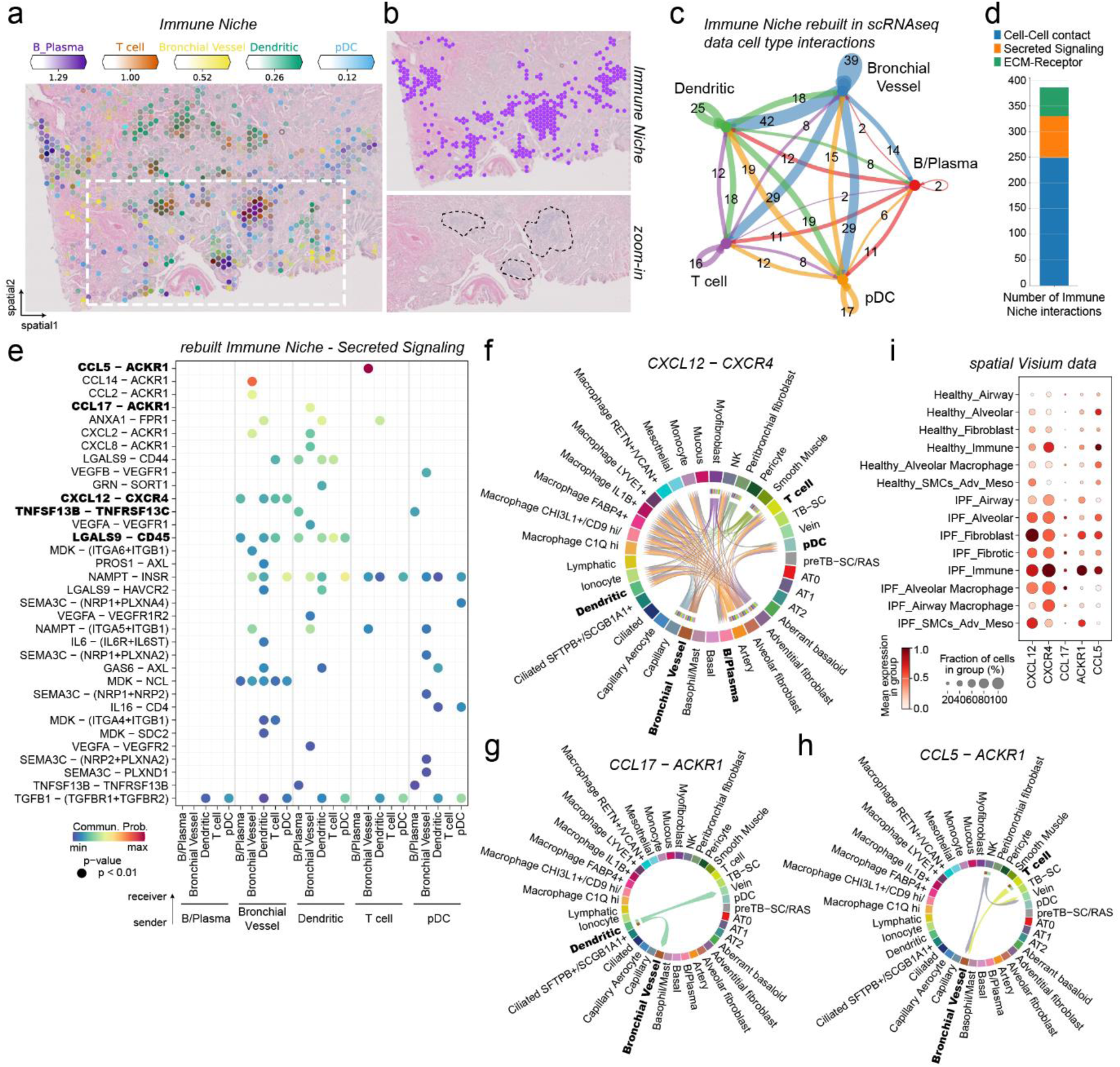
Immune cells from foci and are recruited by IFP specific Bronchial Vessels. **(a-b)** Spatial plots show for one IPF tissue sections **(a)** abundance of Immune niche associated cell types, **(b)** Immune niche distribution as well as zoom into H&E-stained tissue for highlighted regions. **(c)** Bubble plot summarizing the number of interactions between cell types within the rebuilt Immune niche in the scRNA-seq PF-ILD atlas data. **(d)** Number of interactions within the Immune niche per communication category. **(e)** Heatmap of significant ligand-receptor pairs from the Cell-Cell contact category within the Immune niche. **(f-h)** Chord plot visualizing the interacting cell types for the ligand-receptor pair **(f)** CXCL12-CXCR4, **(g)** CCL17-ACKR1 and **(h)** CCL5-ACRK1. **(i)** Gene expression of given receptor and ligand genes across the niches and diagnosis in the spatial Visium data.

We considered how the cells could organize in those foci and analyzed the cell-cell communication by rebuilding the niche again with its contributing cell types in the scRNA-seq dataset (Supplementary table S6). Summarizing the number of all interactions revealed that the Bronchial Vessels seemed at the center of it, contributing most to the ligand-receptor interactions within the niche and with all the other cell types (Fig. 5c).

The Cell-cell communication in the Immune niche was mostly coming from Cell-Cell contact interactions (66%) (Supp. Fig. 15c d), followed by Secreted signaling (Fig. 5e) and ECM-Receptor interactions (Supp. Fig. 15c, e).

Focusing on Secreted signaling, we identified Bronchial Vessels as secreting several chemokines and cytokines able to recruit lymphoid cells (Fig.5e). Bronchial Vessels for example expressed *CXCL12*, also found on Myofibroblast, that is received by the receptor *CXCR4*, found on T cells, B cells, *pDCS* and Dendritic cells (Fig. 5f). But Bronchial Vessels, with their receptor *ACKR1*, seem also to be able to act upon signals via *CCL17* from Dendritic cells or *CCL5* from T cells and Natural killer cells (NK) (Fig. 5g, h). These observations could be validated in the spatial data, with *ACKR1* and *CCXCR4* being most abundant and *CXCL12*, *CCL17* and *CCL5* being highly expressed in the Immune (Fig. 5i).

Overall, the spatial transcriptomics data together with H&E tissue structure, revealed the Immune niche as histopathological feature in IPF, in the form of foci of multiple lymphoid cell types, potentially recruited by distinct secreted signaling of the ectopic endothelial Bronchial Vessels.

## DISCUSSION

Our integration of six already independently published human lung scRNA-seq studies into a harmonized PF-ILD atlas, using state of the art computational tools, was the basis for a robust and generalizable analysis of IPF patients on a single cell level. At the same time, we performed spatial transcriptomic analysis of lung FFPE tissue, comparing for the first time in an unbiased manner and on whole tissue sections, spatially resolved gene expression between IPF and control patients. While the layer of simply visualizing gene expression directly in tissue could already help immensely in understanding of biology and disease, the analysis of in situ cell type distribution provided further in sights. In this study, we mainly made use of the high confidence cell type annotation from scRNA-seq data, that enabled an increased resolution of the spatial transcriptomics data, by estimating the cell-type abundance per spot. Comparing cell type frequencies between the modalities, confirmed a cell type bias in tissue dissociation and sample processing to a single cell solution: fragile epithelial and endothelial cell types were more abundant in the spatial Visium data, while robust myeloid cells such as Macrophages were overrepresented in scRNA-seq data. This bias is also observed when comparing single nuclei data with scRNA-seq data, which also does not require cell isolation, and validates spatial transcriptomics as a tool to capture biology in situ(*14*, *15*, *39*).

We used NMF to detect cellular niches in an unbiased manner across sections and conditions and identified three niches that only exist in fibrotic IPF tissue: the Fibrotic niche, the Airway Macrophage niche, and the Immune niche. Plotting and analyzing the neighborhood of these niches in IPF tissue, offered the picture of a disorganized tissue, compared with control sections. However, we were able to identify a preferential localization of the Fibrotic niche and especially the Aberrant Basaloid cells around airways in the IPF tissue. Aberrant basaloid cells, as a rather novel disease associated cell state of unknown and debated origin, were first identified as distinct from any epithelial cell type due to their mesenchymal and EMT marker expression and association with senescence(*5*, *6*). Here, we demonstrate a co-localization of Aberrant Basaloid cells with Myofibroblasts around airways, that correlated with a senescence gene-signature score.

Cell-cell communication analysis has advanced with available scRNA-seq data and is mostly used as a discovery tool to unveil potentially interacting cell types, based on ligand-receptor pair expression, often without consideration if those cell types are close enough to have a chance to interact(*40*). Leveraging the fact that multiple cells are captured per spot with the Visium technology, we assumed that those cells captured within one spot, would then be close enough contact to interact. Therefore, we focused on these pathological niches and rebuilt them within the scRNA-seq data, based on the spatial information, to analyze their cell-cell communication in a spatially educated manner.

In the Fibrotic niche, Aberrant Basaloid cells and Myofibroblasts, were found to be heavily involved in the Ephrin pathway, based on receptor-tyrosine kinase signaling, so far only associated with proliferative disease such as cancer(*34*). While proliferation in the Fibrotic niche could be present in the form of Fibroblasts, this is in contrast to the senescent feature of Aberrant basaloid cells, calling for further research of the Ephrin pathway and the Fibrotic niche(*29*, *41*, *42*).

The Airway Macrophage niche preferential localized to airways in the IPF tissue. Our analysis revealed that mainly CHIT3L1+ Macrophage co-localize with distal Ciliated cells, TB-SCs in the airway lumen and Aberrant Basaloid cells, while C1Qhi Macrophage additionally appeared in other niches in fibrotic tissue as well. The localization of the CHIT3L1+ Macrophage to the airway lumen challenges the naming convention of airway macrophages but our communication analysis sheds some light on the poorly explored area of the crucial interactions between macrophages and airway epithelium in disease(*43*). The origin of these fibrotic airway associated Macrophages and if varying functions are associated with different localizations need further efforts(*7*). Airway Macrophage and Fibrotic niche both share a similar neighborhood in and around rather large but distal airways of fibrotic lungs, that appear clearly distinct from remodeled small airways in highly fibrotic parenchymal tissue(*9*). The fact, that Aberrant basaloid cells seem to be split between those niches, however, in both cases, localize to and interact with the distal airway epithelium, points towards a potential airway origin of those strongly debated cells(*44*).

We identified the Immune niche as histologically visible foci of immune infiltrates, predominantly consisting of adaptive immune cells, especially B/Plasma and T cells(*45*). Interestingly, those Immune cells co-localized with the recently identified ectopic PLVAP+ endothelial cells. While previous studies report increased abundance in IPF and association to major airways, we find no preferential localization of the Immune niche or Bronchial Vessel(*16*). Our analysis confirmed an increased abundance of Bronchial Vessels, which were better represented in the spatial data than in the scRNA-seq data (Fig. 1c). Of note is their potential ability to recruit the immune cells into those foci, via the CXCL12-CXCR4 axis, thereby including the endothelium into the aberrant remodeling of the distal parenchymal tissue in IPF(*46*).

Our work has several limitations. The spatial transcriptomic analysis, as any omics data, would benefit from an increased sample size that could help to manifest our findings in a broader group of patients of different disease stages, apart from end-stage IPF, as well as a broader selection of tissue locations in the lung. Algorithms to deconvolute data derived from Visium spots into cell type proportions are powerful tools, supported by the nearly whole transcriptome-capturing probe set, but are rely on well-defined and annotated single cell references(*22*, *47*). However, the current resolution of Visium technology with its resolution of 50µm, captures multiple cells per spot, prohibiting direct single cell analysis(*48*). Additionally, the Visium spatial analysis is based on the transcriptomic phenotype of a cell, that can differ from its functional one, defined by protein expression and turnover. It will be interesting to observe, how the upcoming wave of sub cellular resolution techniques(*48*, *49*), yet with only limited number of genes, can supplement our work and move the field even closer to an in situ single-cell analysis in the natural tissue context.

However, despite these limitations, our data provides key starting points for drug discovery in IPF, where beyond the anti-fibrotic standard of care treatments, such as nintedanib(*50*), a profound unmet medical need persists. Traditional tactics to find new targets rely on observing changes in gene expression between healthy and diseased tissue (often bulk RNAseq) followed by in vitro validation experiments which use a single-cell type responding to a target-relevant stimulus. New therapies are needed, that recognize the importance of different disease-associated cell populations beyond isolated myofibroblasts and importantly address the cell-cell miscommunication between those populations, which drives pathology(*51*). Here, we spatially characterize three key disease-specific niches and reveal the lines of communication between cell populations therein. It is from such data, that novel functional in vitro models can be designed - building entirely new test cascades to validate targets which will disrupt disease-driving niches.

## SUPPLEMENTAL FIGURES

**Supplemental Figure 1:**
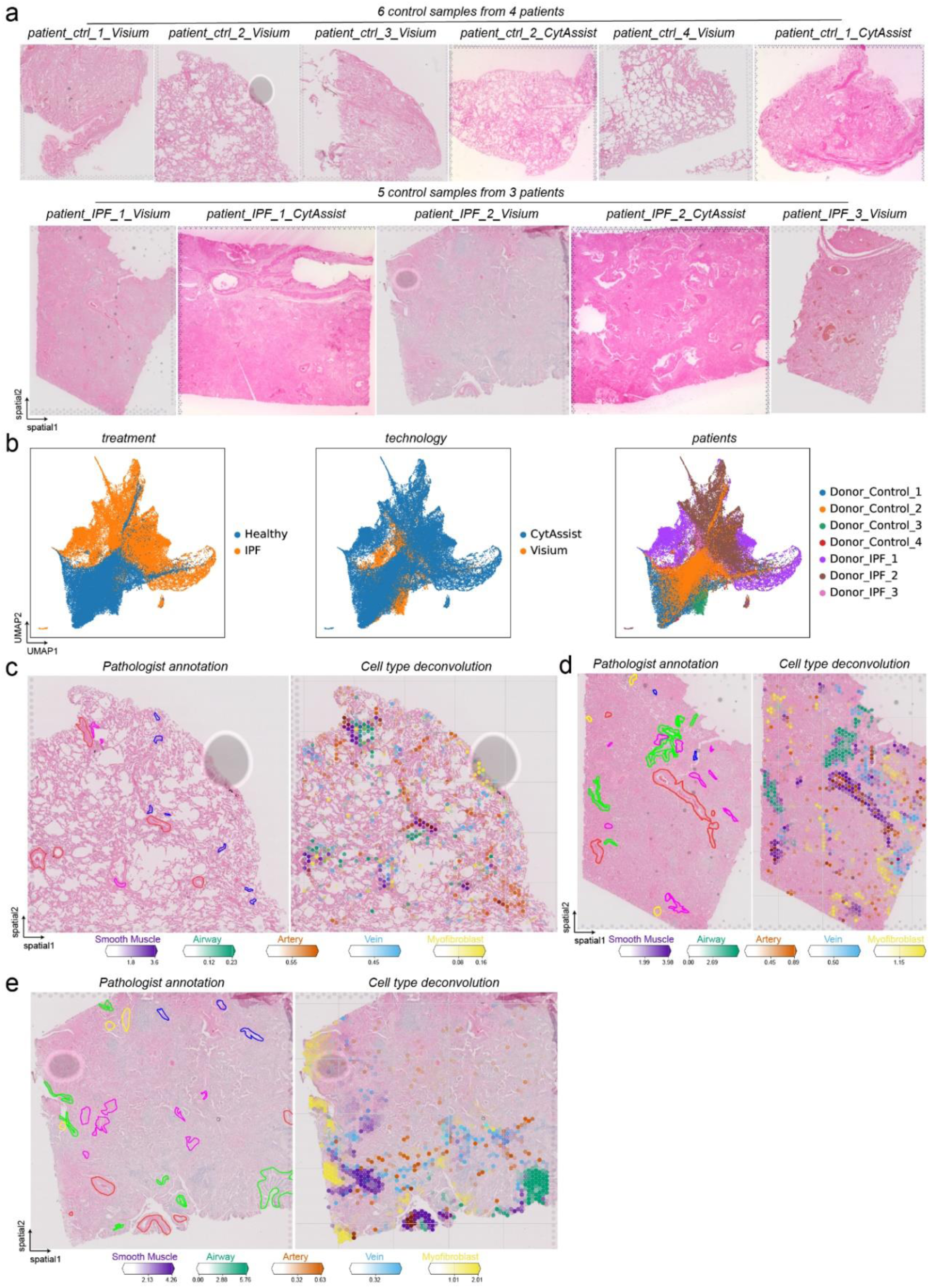
Overview of tissue sections used for Visium for FFPE and CytAssist assays. **(a)** H&E-stained tissue sections of IPF and control patient lungs, used for the indicated assays of either 10x Genomics Visium for FFPE or 10x Genomics CytAssist for FFPE, or both. **(b)** The integrated spatial data of all tissue section is visualized as UMAP. The color code illustrates the disease status, the different technologies, and the donors. **(c-d)** The plots show the comparison of tissue annotated by a pathologist and the respective predicted cell type abundance from cell2location after spot deconvolution for **(c)** a control section and **(d-e)** two IPF sections.

**Supplemental Figure 2:**
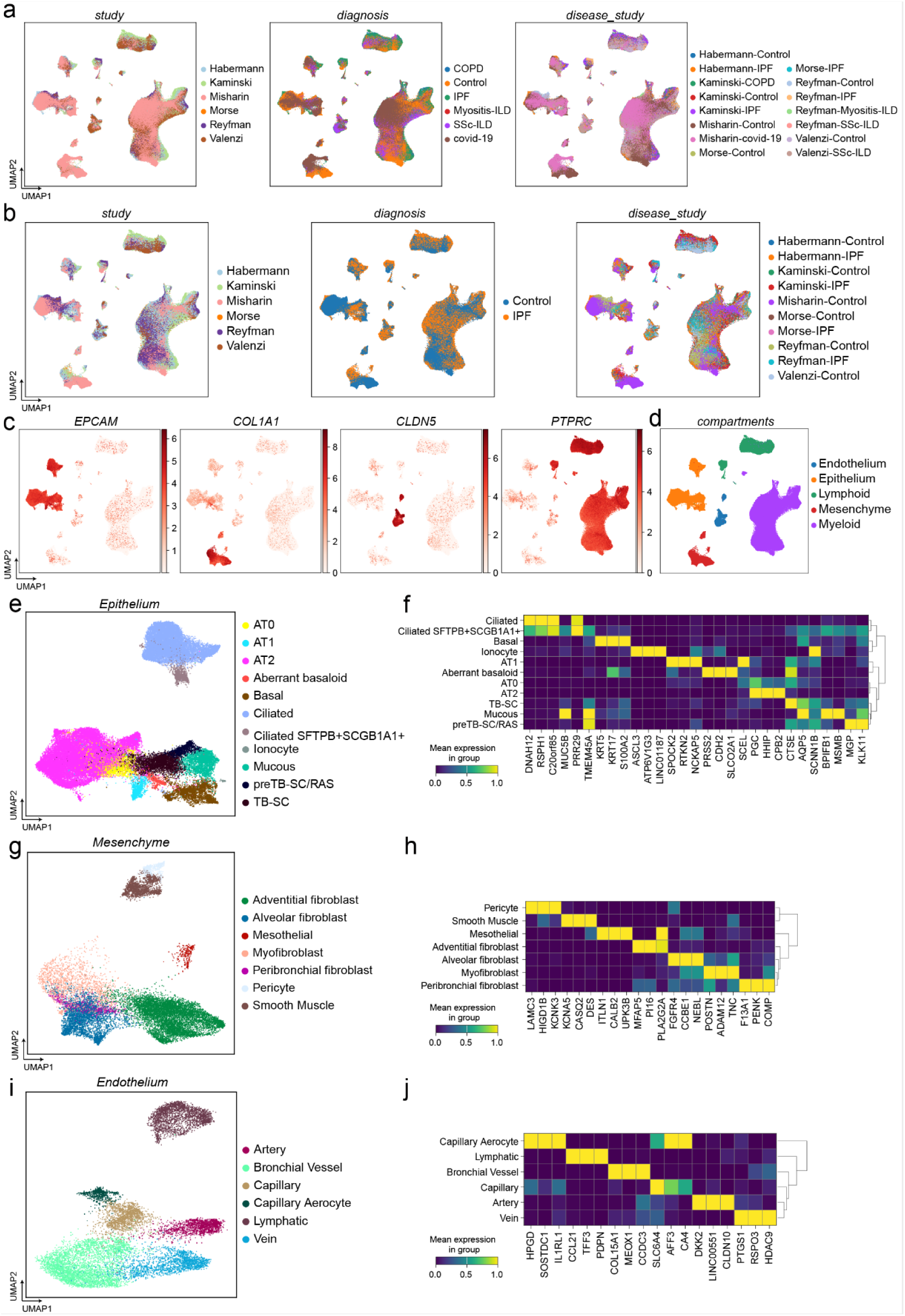
Overview of PF-ILD scRNAseq atlas data and description of compartments. **(a)** The integrated scRNAseq data of the PF-ILD atlas is visualized through Uniform Manifold Approximation and Projection (UMAP). The color code illustrates the different studies/datasets, diagnosis, and their combination. **(b)** UMAPs represent the reduced atlas dataset to only IPF and control diagnosis. The color code illustrates the different studies/datasets, diagnosis, and their combination. **(c-d)** The indicated marker genes were used to select clusters for subsetting the data into compartments. **(e-j)** The compartments are represented with their respective UMAPs and top 2 marker genes per cell type within the compartment, for **(e-f)** EPCAM+ Epithelial cells, **(g-h)** COL1A1+ Mesenchymal cells, **(i-j)** CLDN5+ Endothelial cells.

**Supplemental Figure 3:**
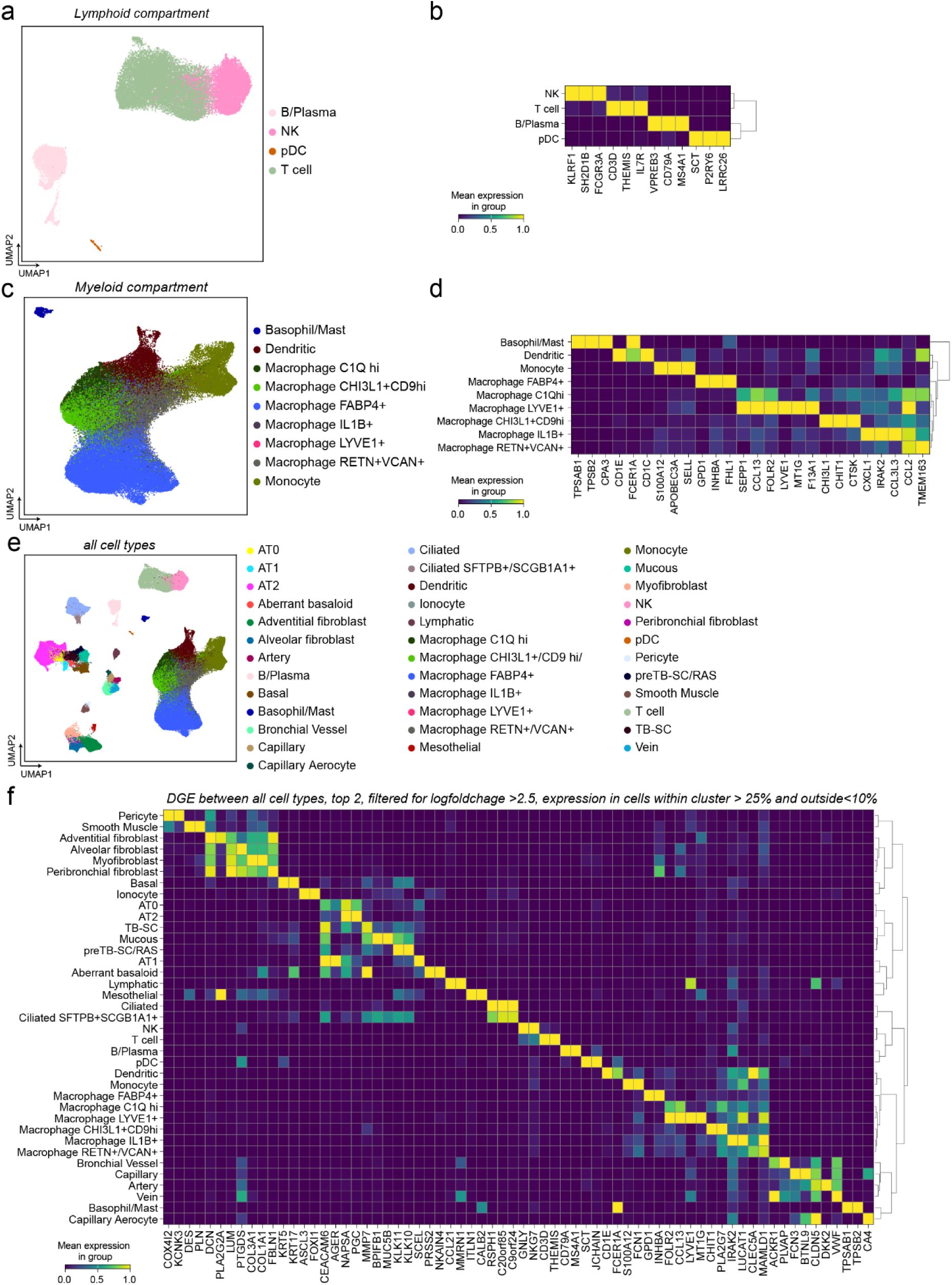
Second part of Overview of scRNAseq atlas data and description of compartments. **(a-d)** The compartments are represented with their respective UMAPs and top 2 marker genes per cell type within the compartment, for **(a-b)** Lymphoid cells, **(c-d)** Myeloid cells. **(e)** The color code of the UMAP indicates all annotated cell types. **(f)** Differential gene expression (DGE) analysis was performed between all cell types and filtered for a log2foldchage > 2.5, expression of the gene in cells within the cluster >25% and in cells outside the cluster < 10%. The heatmap shows normalized mean expression for the Top 2 genes per cell type.

**Supplemental Figure 4:**
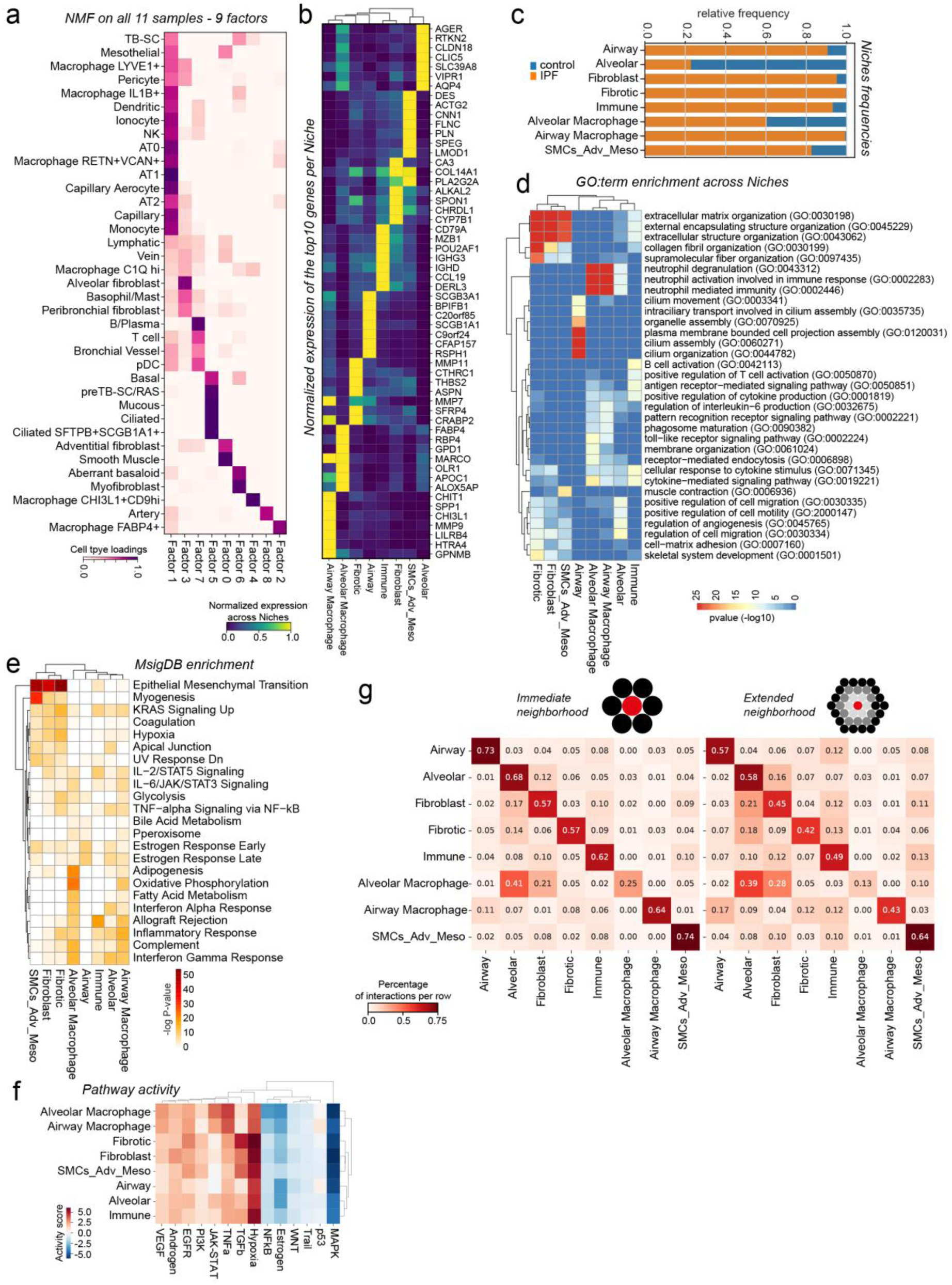
Characterization of NMF inferred cellular niches. **(a)** The heatmap shows normalized cell type abundance for each of the 9 factors as calculated by the non-matrix factorization (NMF) of the cell2location package. Factor 8 only scored for 12 spots and was combined with Factor 0 into the SMCs_Adv_Meso niche. **(b)** The heatmap shows normalized expression of top10 genes per niche as result of the *allmarkers()* function from *scanpy*(***52***). These gene lists sorted by log2foldchange were used for the following enrichment analysis. **(c)** The bar graph illustrated relative frequencies of the niches in IPF or control sections. **(d)** The heatmap shows top3 enriched GO:terms per niche, colored by -log10 pvalue. **(e)** The heatmap shows top enriched Molecular Signature Database (MsigDB) Hallmark genes sets per niche. **(f)** The heatmap shows the pathway activity score per spot, summarized per niche. The R package *decoupleR*(***53***) was used to score the pathway activity as provided by the R package *PROGENy*(***53***). **(g)** The heatmap shows the interactions between the niches as percentage per row, calculated by the function *spatial_neighbors()* in *squidpy*(***54***) for the direct neighbors (1 ring of spots) or the externed neighborhood (3 rings of spots).

**Supplemental Figure 5:**
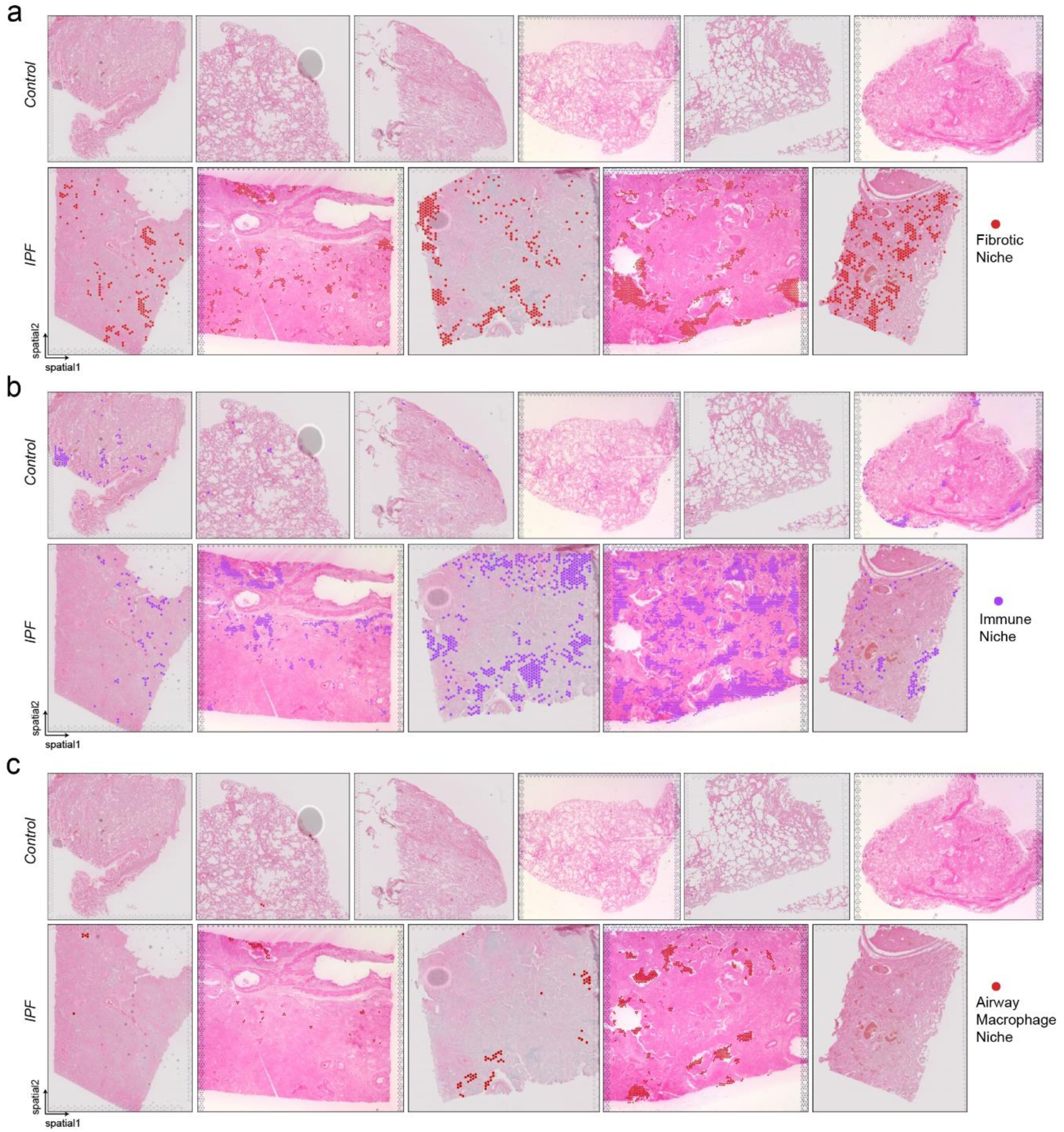
Pathological cellular niches across all tissue sections. **(a-c)** The spatial plots display the distribution of the three in IPF emerging pathological cell type niches across all tissue sections for **(a)** the Fibrotic niche, **(b)** the Immune niche and **(c)** the Airway Macrophage niche.

**Supplemental Figure 6:**
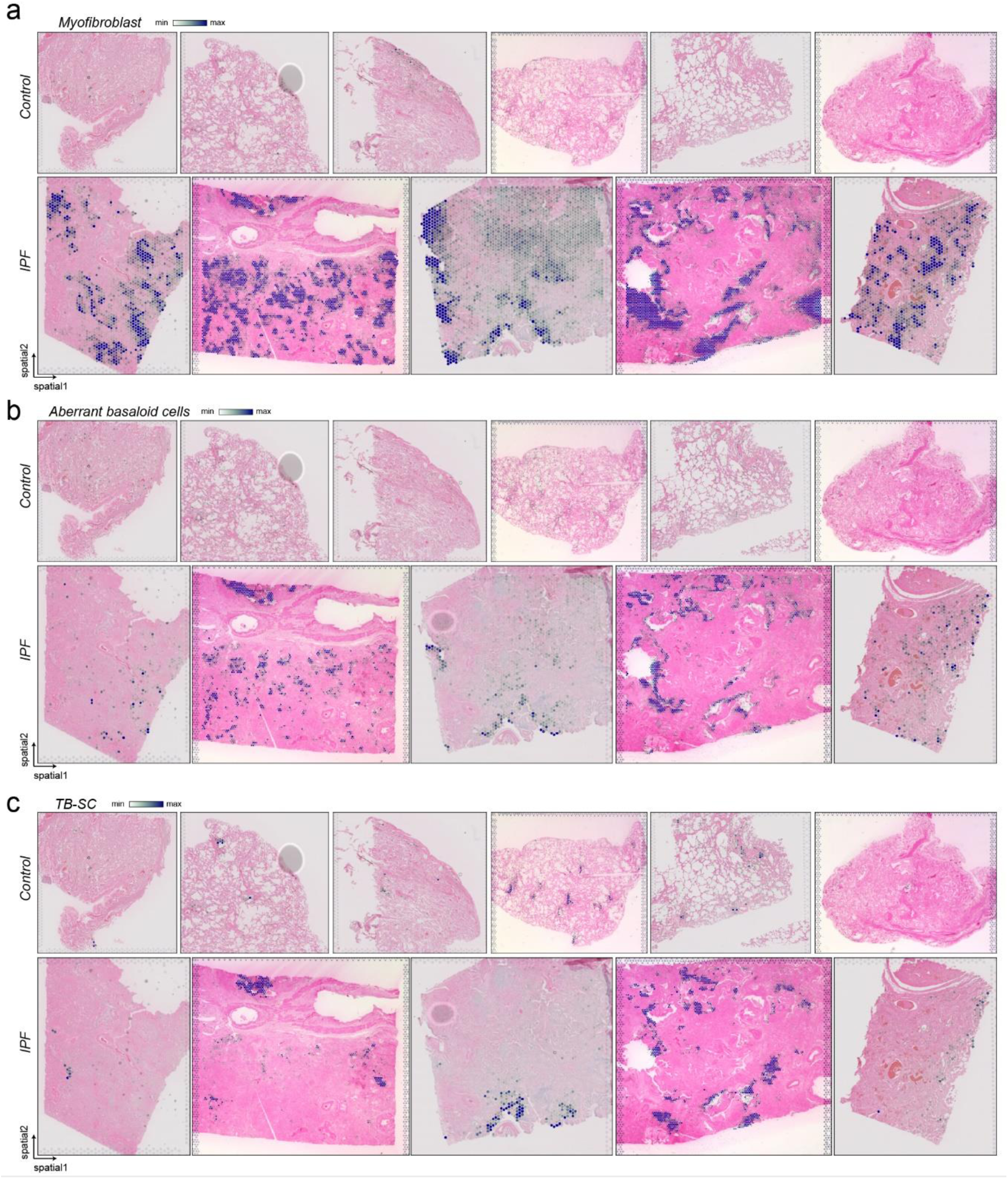
Localization of Fibrotic niche associated cell types. **(a-c)** The spatial plots display cell type abundance across all tissue sections for **(a)** Myofibroblasts, **(b)** Aberrant basaloid cells, and **(c)** TB-SCs.

**Supplemental Figure 7:**
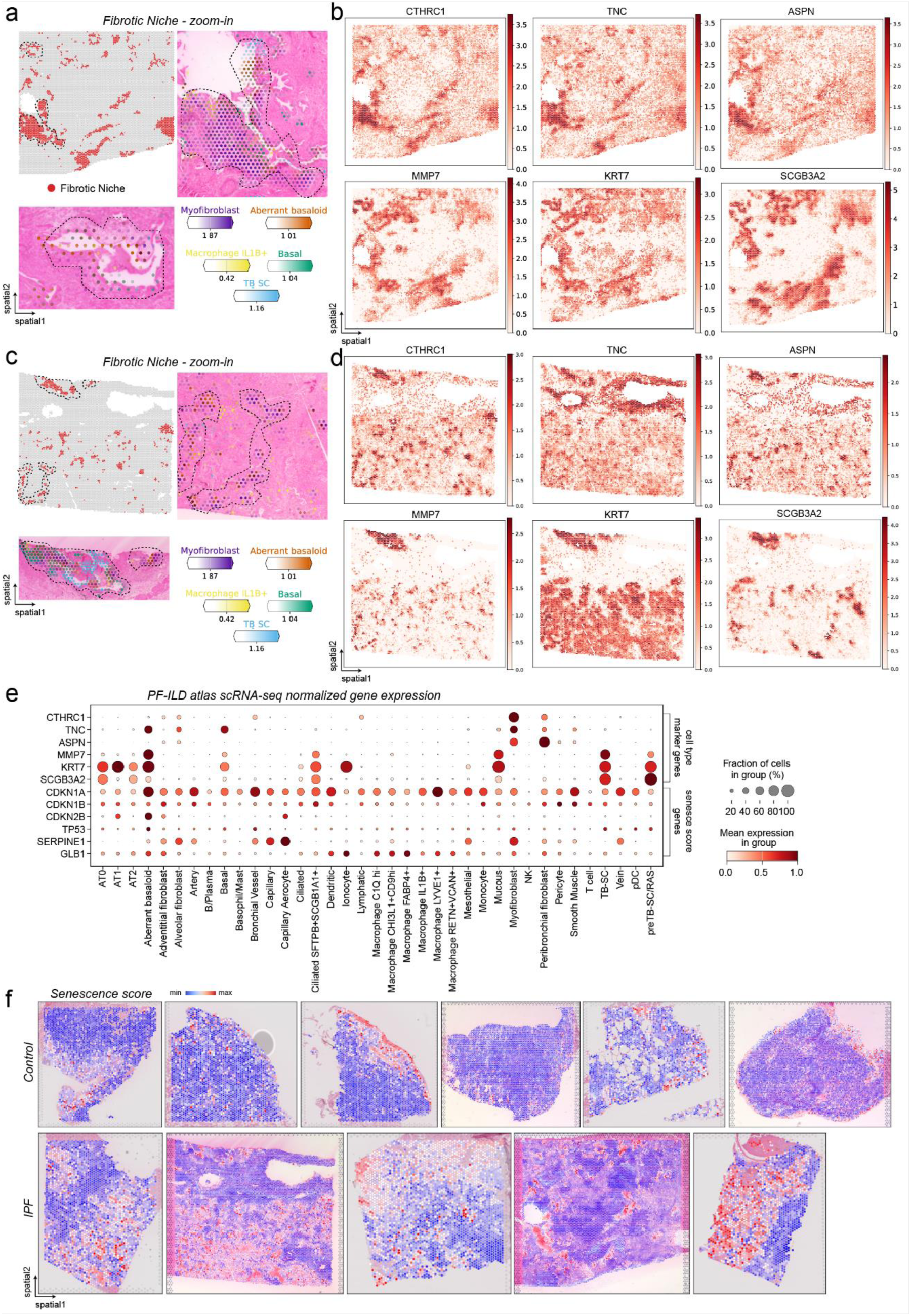
Localization and characterization of Fibrotic niche. **(a-d)** For two IPF tissue sections, the spatial plots display **(a, c)** the distribution of the Fibrotic niche across the slides and zoom ins of two example regions with of selected cell type abundances, as well as **(b, d)** indicated marker gene expression. **(e)** The dotplot displays indicated gene expression across cell types in the IPF single cell atlas. **(f)** Senescence gene score (*CDKN1A*, *CDKN1B*, *CDKN2B*, *TP53*, *SERPINE1*, *GLB1*) calculated using hotspot gene-module scoring(***26***), across all tissue sections.

**Supplemental Figure 8:**
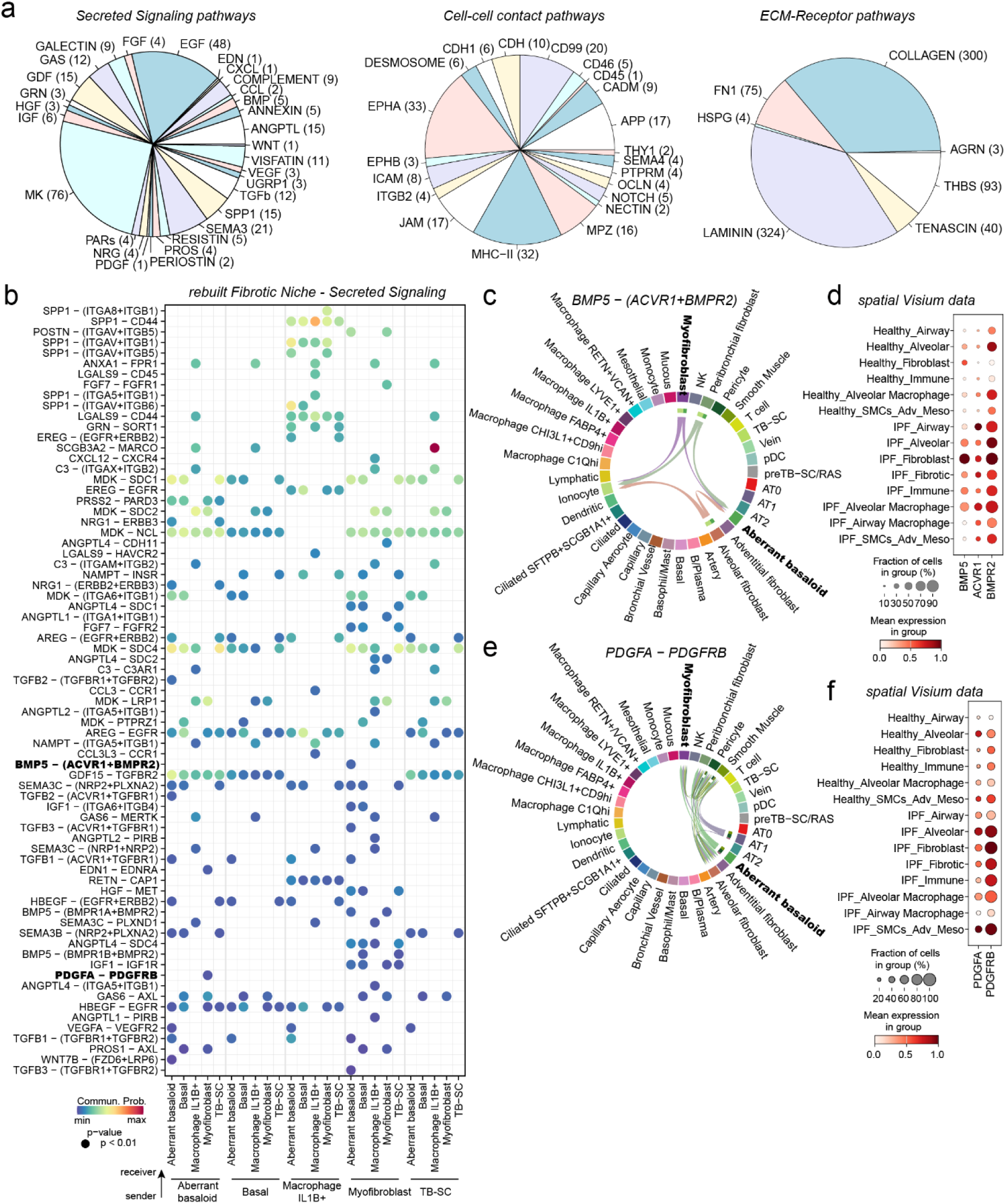
Cell-cell signaling within the Fibrotic niche. **(a)** Pie charts show the distribution of receptor-ligand pairs in Fibrotic niche into the three cell communication categories and their associated signaling pathways. **(b)** Heatmap of significant ligand-receptor pairs from the Secreted Signaling category within the Fibrotic niche. **(c-f)** Chord plot visualizing the interacting cell types and heatmaps the expression of the involved genes in the spatial data for the ligand-receptor pairs **(c, d)** *BMP5*-(*ACVR1*+*BMPR2*) and **(e, f)** *PDGFA*-*PDGFRB*).

**Supplemental Figure 9:**
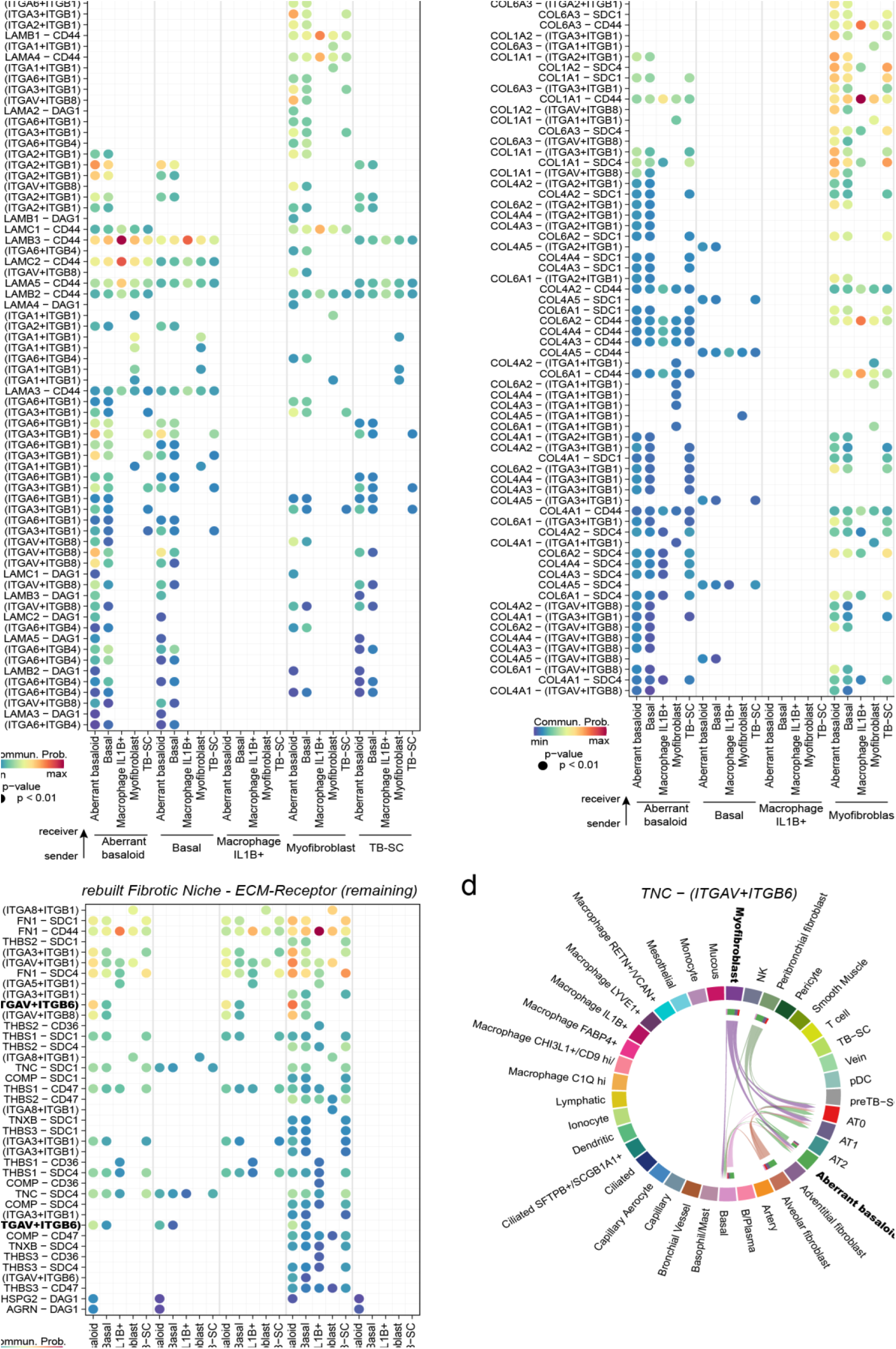
ECM-Receptor signaling within Fibrotic niche. **(a-c)** Heatmaps of significant ligand-receptor pairs from the ECM-Receptor category within the Airway Fibrotic niche, split into major pathways **(a)** Laminin pairs, **(b)** Collagen pairs and **(c)** the remaining pairs. **(d)** Chord plot visualizing the interacting cell types for the ligand-receptor pair *TNC*-(*ITGAV*+*ITGB6*).

**Supplemental Figure 10:**
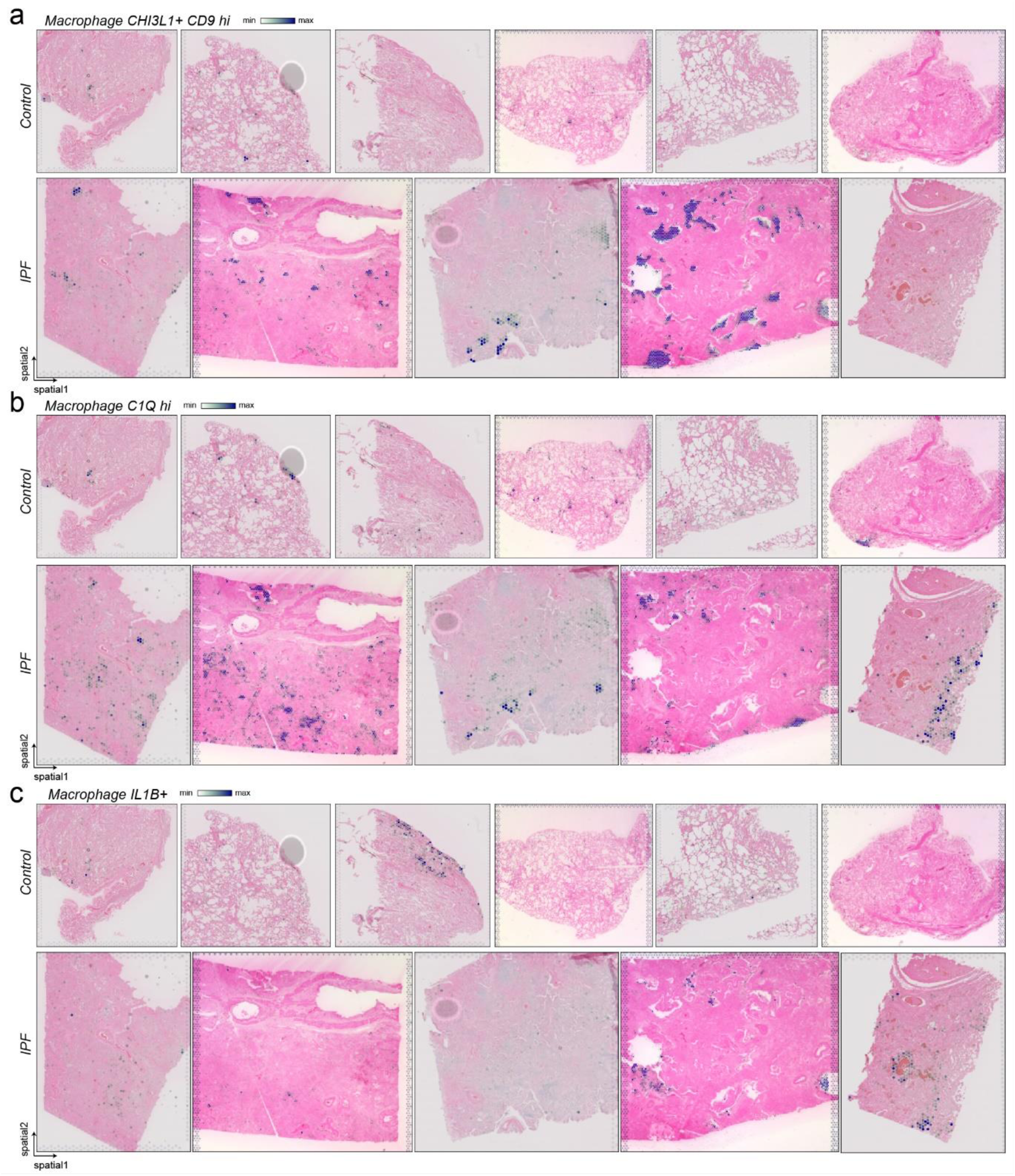
Localization of disease induced Macrophage cell types. **(a-c)** The spatial plots display cell type abundance across all tissue sections for the different Macrophage cell subtypes of **(a)** CHI3L1+ CD9hi Macrophages, **(b)** C1Qhi Macrophages and **(c)** IL1B+ Macrophages.

**Supplemental Figure 11:**
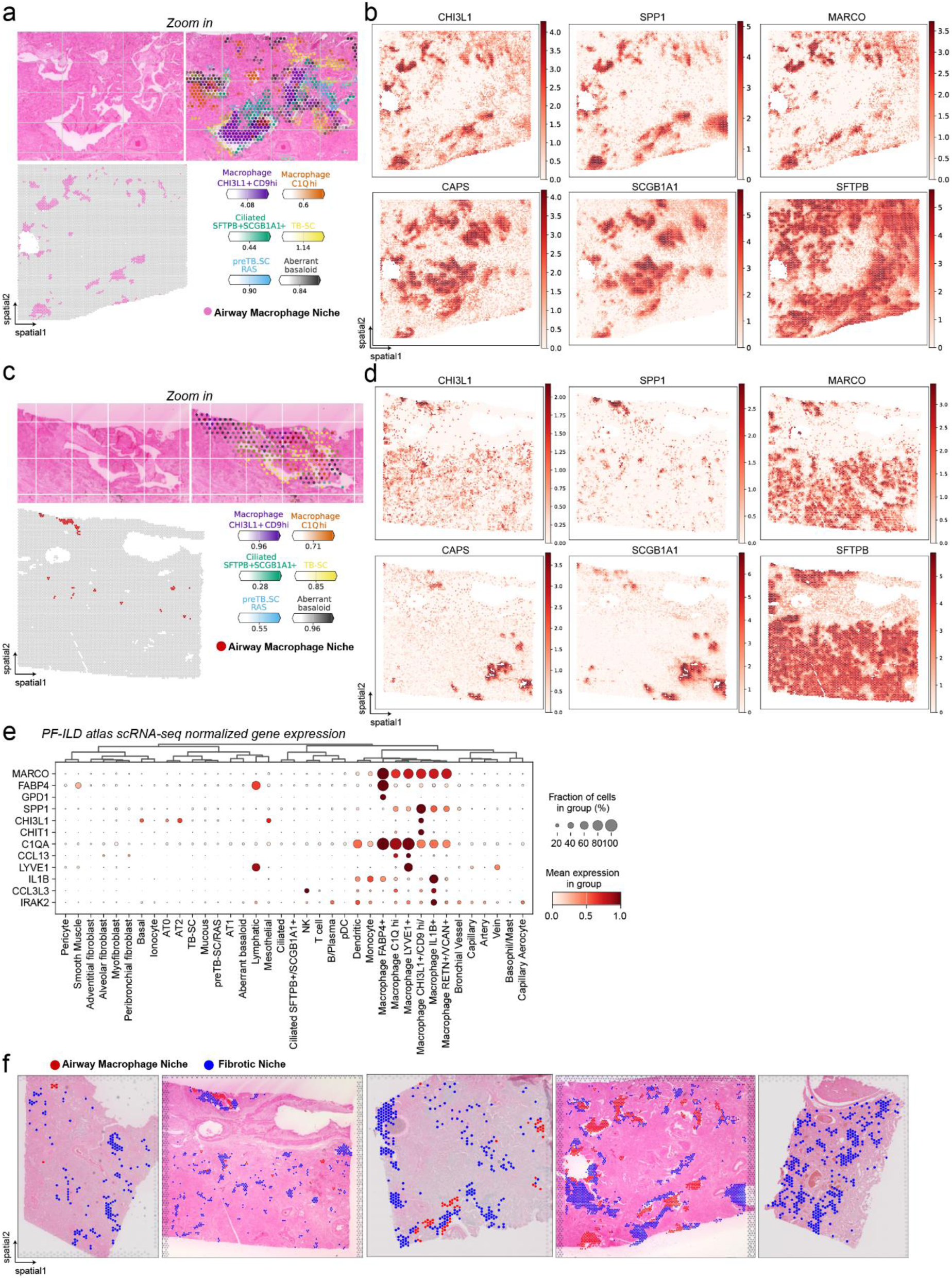
Localization and characterization of the Airway Macrophage niche. **(a-d)** For two IPF tissue sections, the spatial plots display **(a, c)** the distribution of the Airway Macrophage niche across the slides and zoom ins of two example regions with of selected cell type abundances, as well as **(b, d)** indicated marker gene expression. **(e)** The dotplot displays indicated gene expression across cell types in the IPF single cell atlas. **(f)** The spatial plots show localization of the Airway Macrophage niche and the Fibrotic niche across the 5 IPF tissue sections.

**Supplemental Figure 12:**
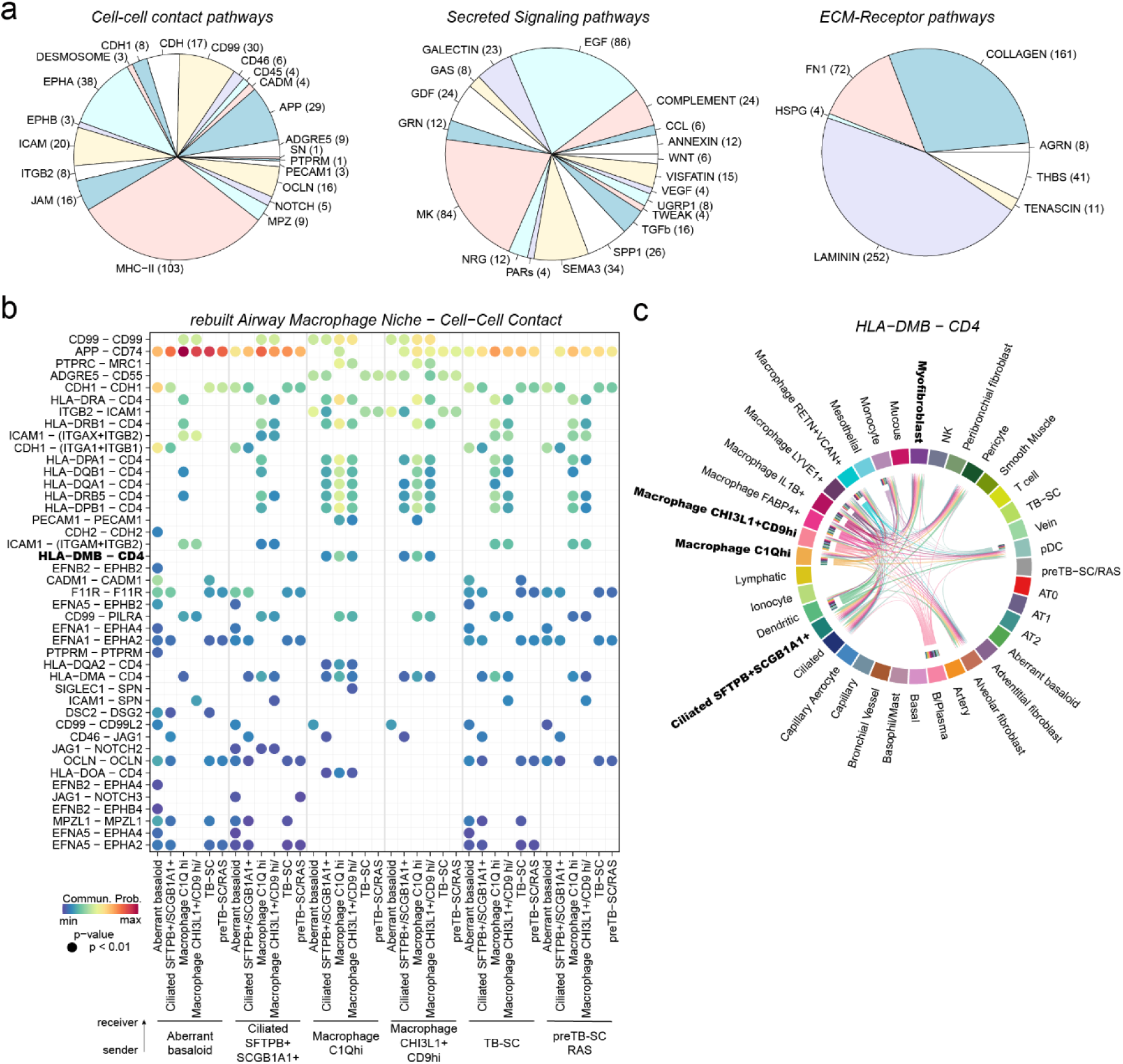
Cell-cell signaling within the Airway Macrophage niche. **(a)** Pie charts show the distribution of receptor-ligand pairs in the Airway Macrophage niche into the three cell communication categories and their associated signaling pathways. **(b)** Heatmap of significant ligand-receptor pairs from the Cell-Cell contact category within the Airway Macrophage niche. **(c-f)** Chord plot visualizing the interacting cell types and heatmaps the expression of the involved genes in the spatial data for the ligand-receptor pair HLA-DMB – CD4.

**Supplemental Figure 13:**
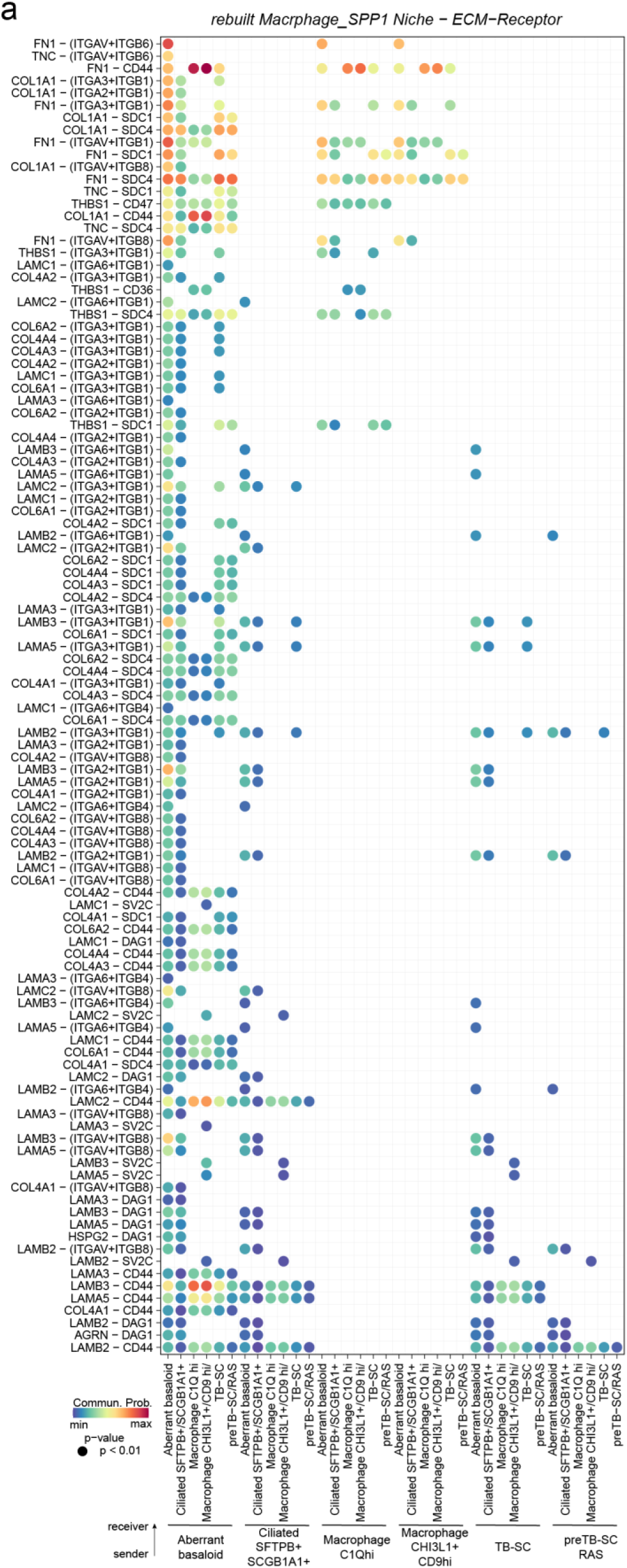
ECM-Receptor signaling within Airway Macrophage niche. **(a)** Heatmap of significant ligand-receptor pairs from the ECM-Receptor category within the Airway Macrophage niche.

**Supplemental Figure 14:**
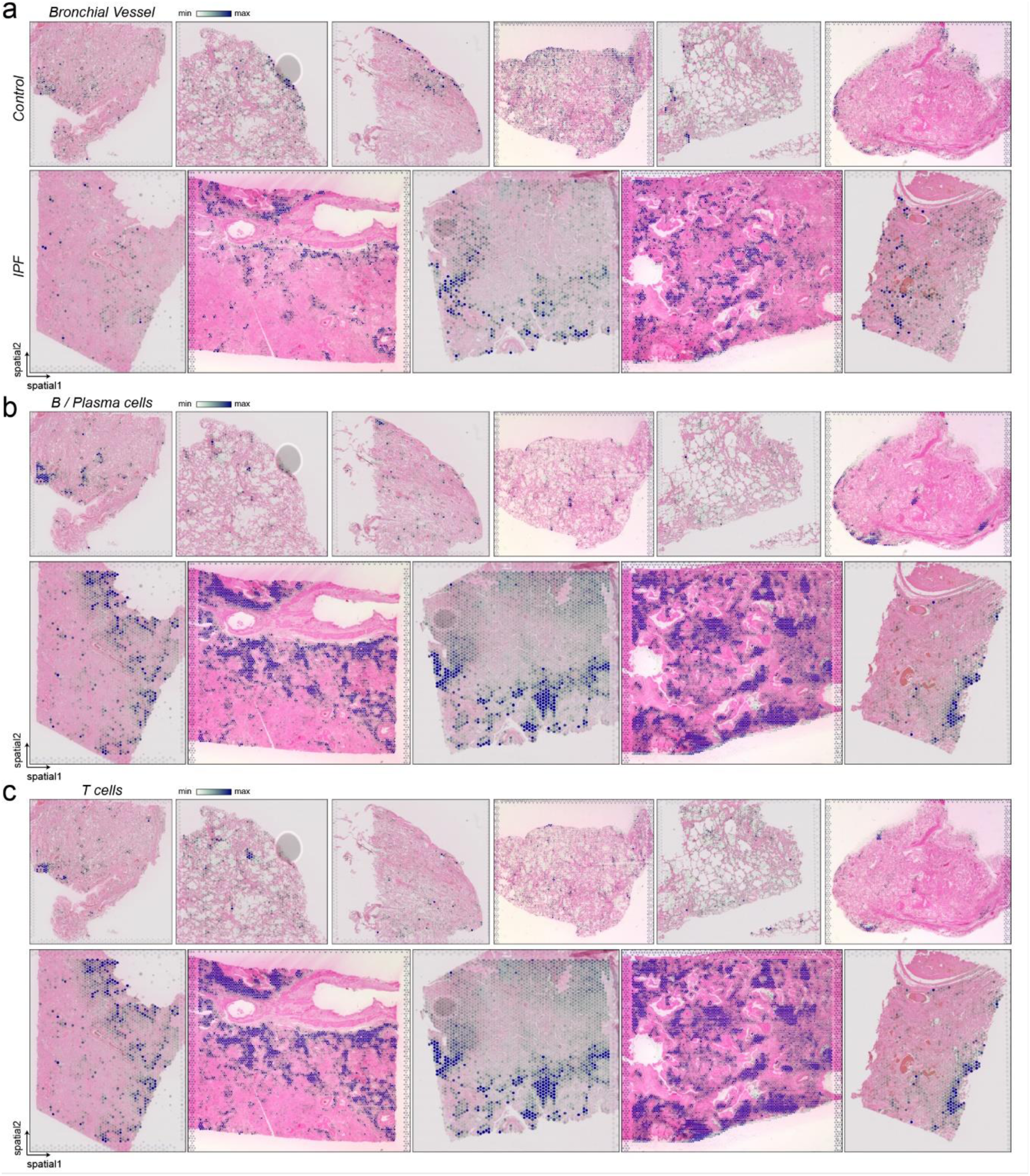
Localization of Immune niche contributing cell types. **(a-c)** The spatial plots display the cell type abundance for indicated immune cell types across all tissue sections for **(a)** Bronchial Vessel cells, **(b)** B and Plasma cells and **(c)** T cells.

**Supplemental Figure 15:**
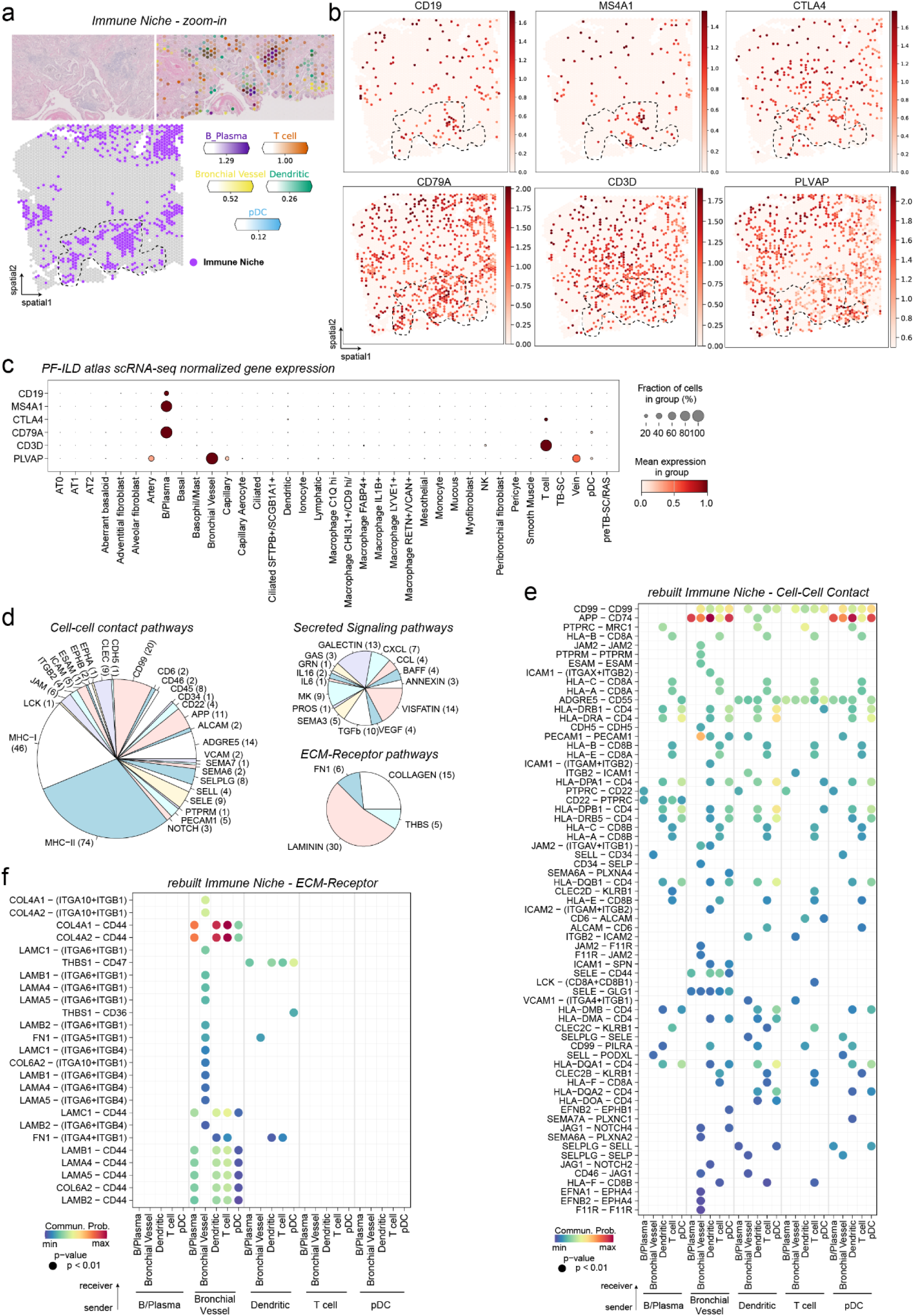
characterization of the Immune niche. **(a-b)** For one IPF tissue section, the spatial plots display (a) the distribution of the Fibrotic niche across the slides and zoom ins of two example regions with of selected cell type abundances, as well as (b) indicated marker gene expression. **(c)** The dotplot displays indicated gene expression across cell types in the IPF single cell atlas. **(d)** Pie charts show the distribution of receptor-ligand pairs in the three cell communication categories and their associated signaling pathways. **(e-f)** Heatmaps of significant ligand-receptor pairs from the **(e)** Cell-Cell Contact and **(f)** the EMC-Receptor category within the Immune niche.

## SUPPLEMENTARY TABLES

**Supplementary table S1:** Marker genes scRNA-seq data (cell type level)

**Supplementary table S2:** Baseline characteristics of human IPF and control samples used for Visium

**Supplementary table S3:** Marker genes Visium data (niche level)

**Supplementary table S4:** Cell-cell communication results with receptor-ligand pairs from CellChat on the rebuilt Fibrotic niche in IPF scRNA-seq data

**Supplementary table S5:** Cell-cell communication results with receptor-ligand pairs from CellChat on the rebuilt Macrophage SPP1 niche in IPF scRNA-seq data

**Supplementary table S6:** Cell-cell communication results with receptor-ligand pairs from CellChat on the rebuilt Immune niche in IPF scRNA-seq data

**Supplementary table S7:** Makers used to annotate cell types in the integrated PF-ILD scRNA-seq atlas.

## ACKNOWLEDGMENTS

We thank all the patients and their families for supporting the progress of science. We thank Diana Reinhardt for assistance in FFPE tissue sectioning and Werner Rust for technical assistance in next-generation sequencing. We are grateful to Jakub Widawski for the help in integrating the PF-ILD atlas.

## AUTHOR CONTRIBUTIONS

MJT, FR and CHM conceived and designed the study and MJT and FR supervised the entire work. CHM wrote the paper. CHM generated custom code and performed bioinformatic analysis on the scRNA-seq and Visium data. DS performed Visium experiments. SJ sectioned, stained, and imaged the tissue sections. JD supervised tissue work. AD supervised Visium and sequencing work. CL performed pathological tissue annotation. LN, BR, JK sourced tissue. DJ and MK provided resources. MJT, FR, HS and AD provided resources and funding. All authors read and corrected the final manuscript.

## COMPETING INTERESTS

CHM, DS, JS, CL, HS, JD, AD, FR and MJT are current employees of Boehringer Ingelheim. All other authors declare that they have no competing interests relating to this work.

## MATERIALS AND METHODS

### Data availability

Raw 10X Genomics Visium data generated for this study will be made available on GEO upon publication. Processed count tables of the Visium data (10.5281/zenodo.10012934) and the integrated PF-ILD atlas (10.5281/zenodo.10015169) are available on Zenodo. The publicly available human lung IPF single-cell RNA-seq datasets used to integrate the PF-ILD atlas can be accessed under the following GEO accession numbers: GSE128169 (Valenzi)(*8*), GSE128033 (Morse)(*7*), GSE135893 (Habermann)(*5*), GSE122960 (Reyfman)(*4*), GSE136831 (Kaminski)(*6*), GSE158127 (Misharin)(*21*).

### Code availability

Code and Jupyter notebooks that were used to analyze the are deposited at GitHub and Zenodo: https://github.com/christophhmayr/2023_Mayr

### Integration of public scRNA-seq datasets into PF-ILD atlas

Single cell count matrices containing interstitial lung diseases and control samples were identified and downloaded from GEO. Samples were processed following best practices(*55*) using scanpy(*52*) and integrated using scVI(*56*). For integration of data we used the union of all common genes between the datasets and then selected 6500 highly variable genes using the Seurat v3 method. The individual single cell samples were used for batch correction and as co-variates we used the percentage of mitochondrial genes and the lab that generated the data (eg. Adams. Et al dataset). While mitochondrial percentage is a common covariate used, we reasoned that batch effects from each sample and from the lab differences in tissue dissociation and processing steps in the lab and in-silico should be considered. The parameters used were as follow: n_hidden: 128, n_latent: 10, n_layers: 1, dropout_rate: 0.1. The integrated dataset contains 405.741 cells from 155 samples for the following diagnosis: IPF (n=65), COPD (n=17), SSc-ILD (n=12), Myositosis-ILD (n=1), Hypersensitivity pneumonia (n=1), Covid (n=3), Control (n=57). For analysis in this manuscript, an IPF version of the atlas was used, subsetting the PF-ILD atlas by diagnosis to only IPF and control patients, as well as downsampling the number of cells to max 1000 per sample.

### Cell type annotation

Cell lineages on the dataset were identified using broad clustering in combination with well-known markers as follows: Epithelial (EPCAM), Endothelial (CLDN5), Mesenchyme (COL1A1), and Immune (CD45 – official gene name: PTPRC) which we subdivide into Myeloid (FCER1G) and Lymphoid (FCER1G negative). For further annotations the cell lineages were analyzed individually, based on well-known markers as well as the markers from the original dataset publications (for markers used to annotate cell types see Suppl. Table S7)(*57–62*).

### Human tissue and ethics statement

Human FFPE tissue was sourced from a commercial vendor (BioIVT). Fresh human lung tissue from explants and resections was collected at Hannover Medical School. All organs were picked up at the operation theatre immediately after surgery and worked up using standardized protocols as described elsewhere(*63*). Briefly summarized, obtained samples were fixed in 3.7% formaldehyde, subsequently dehydrated, and eventually embedded in paraffin by means of an automatic tissue-processing device (LOGOS microwave hybrid tissue processor, Milestone Medical, Sorisole, Italy). All patients gave written informed consent. The study was approved by and conducted according to the requirements of the local ethics committee at Hannover Medical School (8867_BO_K_2020).

### Tissue preparation for spatial transcriptomics

Sectioning: Tissue sections were prepared according to the Visium Spatial Gene Expression for FFPE (CG000408) or Visium CytAssist Tissue preparation guide (CG000518). Briefly, after cooling and re-hydration of tissue blocks in an ice bath, 5µm sections were prepared and transferred to a RNAse-free water bath at 42 °C. Sections were placed on either Visium Spatial Gene Expression Slides directly (10x Genomics, standard workflow) or SuperFrost Plus microscopy slides (CytAssist workflow) and dried for 3h in an oven at 42°C, followed by overnight drying at room temperature. Staining: Deparaffinization and H&E staining was performed according to Visium user guides (CG000409, CG000520). Briefly, sections were incubated for 2h at 60°C in a dry oven, before deparaffinization in Xylene and rehydration using an ethanol gradient were performed. Sections were mounted using 85% glycerol for the H&E imaging step. Imaging: H&E-stained slides were scanned on a Leica Axioscan brightfield scanner using 20x magnification. Coverslips were removed by immersing the mounted slides in Milli-Q water until the coverslip detaches. Annotation: Using the freehand annotation function in HALO Link (IndicaLabs), examples of distinct histological structures (arteries, veins, airways, fibroblastic foci, muscle) were annotated by a pathologist.

### Visium spatial transcriptomics and sequencing

Following imaging, 11 H&E-stained tissue sections from three IPF and three control patients were processed with the standard Visium spatial for FFPE Gene expression kit, Human transcriptome (7 sections, 10x Genomics Cat #1000334) or the Visium CytAssist Spatial Gene Expression for FFPE, Human transcriptome, 11mm (4 sections, 10x Genomics Cat #1000444) as per manufacturer’s instructions (CG0000407, Rev D; CytAssist CG000495, Rev C). Briefly, deparaffinized and decrosslinked H&E-stained sections were hybridized with Human WT probes (10x Genomics) directly on visium slides (standard) or on SuperFrost Plus microscopy slides (CytAssist) on a thermal cycler at 50°C for ∼18hrs followed by probe ligation, CytAssist-enabled RNA degradation and tissue removal (CytAssist workflow only) extension, elution and library preparation as per manufacturer’s instructions. Visium libraries were analyzed via qPCR (QuantStudio 6, Applied Biosystems) and a total of 18-20 (standard workflow) or 12-14 (CytAssist workflow) PCR cycles were employed for library amplification. Visium libraries were quantified with Qubit 1x dsDNA HS Assay Kit (Invitrogen, Cat # Q33231) on a Qubit™ 4 Fluorometer (Invitrogen). Qualitative assessment of Visium libraries was conducted with the High Sensitivity NGS Fragment Analysis Kit (Agilent, DNF-474) on a 96-channel Fragment Analyzer (Agilent) to assess size distribution and adapter dimer presence (<0.5%). All Visium libraries were normalized, pooled and spiked with 10% PhiX Control v3 (Illumina) then loaded at 0.75nM and sequenced paired-end (Rd1:28 Rd2:10 Rd3:10 Rd4:50) on a Novaseq 6000 (Illumina).

### Preprocessing of spatial Visium RNA-seq data

The Spaceranger computational pipeline from 10x Genomics was used to generate count matrices (v1.3.1 and v2.0 for CytAssist samples). The reads were aligned to the hg38 human reference genome (GRCh38-2020-A) and against the probe set references for human provided by 10x Genomics (v1 for FFPE samples and v2 for CytAssist samples). After preprocessing, analysis of the spatial Visium RNA-seq data was performed with the Python packages scanpy (version 1.9.3)(*52*), squidpy (1.2.3)(*54*, *64*) and cell2location (0.1.3)(*65*). The Spaceranger output files and the corresponding histology images were merged into a single anndata(*66*) object. QC parameters were assessed, and the data filtered to remove spots with less than 800 counts, more than 45000 counts, less than 800 genes, or fewer than 25 cells. We log transformed the data via Scanpy’s pp.log1p() function and calculated highly-variable genes (HVGs) (top 6000 genes with flavor = seurat_v3). Using the HVGs as input we employed scvi-tools(*56*, *67*) to model the latent variables that were then used for UMAP and Leiden clustering, using 30 neighbors, which then was both transferred back to the full-gene dataset.

### Visium spot cell type deconvolution with cell2location and niche identification using NMF

For Visium spot deconvolution into cell types, we used cell2location and followed the tutorial provided on their github(*65*). We loaded the PF-ILD atlas, filtered for IPF and control diagnosis, and performed permissive gene selection with standard settings, before estimating reference cell type signatures with batch_key set to sampleID and the model training set to 250 epochs. Next, both the reference and the spatial input data were subsetted to shared genes, the cells per location parameter set to 8 and detection alpha to 20. The cell2location model was trained with 15000 epochs and the 5% quantile of the estimated posterior distributions of cell abundance exported to the anndata object.

To identify cellular niches across tissue sections, we used the NMF function of the cell2location package, on the predicted cell type abundances. We calculated 3 to 30 factors and found 9 factors to best represent cellular niches, with more factors only splitting the data into single cell types. With the factor for Artery only scoring for a handful of cells, we combined it with the SMCs_Adv_Meso factors to end up with 8 niches.

### Cell type frequency comparison

To compare the cell type frequencies between the modalities, we averaged the normalized cell type frequency per cell in the scRNA-seq data, and the normalized estimated cell type frequency per spot in the spatial Visium data, across the samples and diagnosis.

### Gene pathway activity scoring

For each spot, we estimated signaling pathway activities with PROGENy’s (v1.12.0) weighted human model matrix using signature genes with a p-value cutoff of 0.01. To infer pathway enrichment scores per spot, we ran the Multivariate Linear Model (mlm) from decoupleR(*53*) and summarized the scores across niches.

### Neighbor interactions

We first calculated a connectivity graph using the squidpy’s gr.spatial_neighbors() method. This graph consists of cells (nodes) and cell-cell interactions (edges). Next, we wanted to identify niches that are spatially enriched for one another using a neighborhood enrichment test gr.nhood_enrichment(). Accessing the edges count, we normalized the interactions per niche and compared them between diagnosis, by manually adding those niches that only exist in of the diagnosis. We extended this analysis, comparing immediate and extended niche neighborhood, by calculating the connectivity graphs for 1- or 3-rings of spots around a center target spot.

### GO and MsigDB enrichment

We used scanpy’s tl.rank_genes_groups() to calculate niche specific marker genes using the wilcoxcon test, and filtered the results for genes to have a adjusted pvalue smaller than 5%, a log2foldchange greater than 0.5, be expressed within more than 25% of spots of a niche and less 25% outside the niche. Gene Set Enrichment Analysis was then performed using GSEApy(*68*) (1.0.0) with the libraries “GO_Biological_Process_2021” and “MSigDB_Hallmark_2020” from the human database. Gene sets were considered enriched in the respective signature at a statistically significant level if the FDR-corrected p-value was below 0.01. Signatures were plotted in the heatmap if they were significant in at least one niche.

### Gene signature scoring using hotspot

For scoring the senescence gene signature of the genes GLB1, TP53, SERPINE1, CDKN1A, CDKN1B, CDKN2B, we used the compute_scores function from hotspot(*69*). In short, the counts for each gene are fit into the DANB model (depth adjusted negative binomial), the values for each gene are then scaled (centered), then smoothed based on neighbors of each cell. Finally, these values for all genes in the signature are dimensionality reduced using PCA. We used the log1p counts layer, 30 neighbors, and the X_scVI embedding to calculate the score.

### Cell-cell communication analysis of rebuilt spatial niches in scRNA-seq data

To analyze the spatially informed rebuilt cellular niches in the scRNA-seq data we used CellChat(*70*). We created the CellChat database with the human interactions The cellchat objects were created from the PF-ILDatlas subsetted to only the IPF and control diagnosis samples, and were run separately per condition, before being combined for the analysis. The niches were rebuilt, by selecting the cell types that were overrepresented in the NMF derived niches/factors (Fig. 2a), and subsetting the CellChat object accordingly. The interactions per niche were filtered for a p-value smaller than 0.01. For a better representation we split the analysis and figure panels based on the CellChatDB annotation categories that group the interactions into Cell-Cell Contact, Secreted Signaling and ECM-Receptor.

